# The proteasomal deubiquitinating enzyme PSMD14 regulates macroautophagy by controlling Golgi-to-ER retrograde transport

**DOI:** 10.1101/2020.01.29.925503

**Authors:** HA Bustamante, K Cereceda, AE González, GE Valenzuela, Y Cheuquemilla, S Hernández, E Arias-Muñoz, C Cerda-Troncoso, S Bandau, A Soza, G Kausel, B Kerr, GA Mardones, J Cancino, RT Hay, A Rojas-Fernandez, PV Burgos

## Abstract

Ubiquitination regulates several biological processes. Here, we search for ubiquitin-related genes implicated in protein membrane trafficking performing a High-Content siRNA Screening including 1,187 genes of the human “ubiquitinome” using Amyloid Precursor Protein (APP) as a reporter. We identified the deubiquitinating enzyme PSMD14, a subunit of the 19S regulatory particle of the proteasome, specific for K63-Ub chains in cells, as a novel key regulator of Golgi-to-endoplasmic reticulum (ER) retrograde transport. Silencing or pharmacological inhibition of PSMD14 caused a robust and rapid inhibition of Golgi-to-ER retrograde transport which leads to a potent blockage of macroautophagy by a mechanism associated with the retention of Atg9A and Rab1A at the Golgi apparatus. Because pharmacological inhibition of the proteolytic core of the 20S proteasome did not recapitulate these effects, we concluded that PSMD14, and their K-63-Ub chains, act as a crucial regulator factor for macroautophagy by controlling Golgi-to-ER retrograde transport.

## Introduction

Several post-translational modifications (PTMs) contribute to membrane transport ^1, 2^. Among these, ubiquitination acts as a relevant player in protein membrane trafficking ^3–5^ participating as an important protein localization signal ^3, 6–8^. However, to date most studies have focused on its role in endocytosis, particularly in the inclusion of cargoes in intraluminal vesicles (ILVs) of multivesicular bodies (MVBs) ^9–11^ such as cell-surface receptors ^12^. Ubiquitination is a regulated post-translational modification that conjugates ubiquitin (Ub) to lysine (K) residues and is involved in many cellular pathways including the degradation of target proteins through the proteasomes and autophagy ^13, 14^. Ubiquitin modification is a cascade of reactions catalyzed by three classes of enzymes (E1, E2 and E3). An ATP dependent E1 activating enzymes first forms a covalent intermediate with ubiquitin, followed by the transfer of Ub to E2 conjugating enzymes and finally the covalent attachment of Ub to a K residue in the target protein by E3 Ub ligases ^15^. Ubiquitination is a reversible reaction, with specific deubiquitinating enzymes (DUBs) that catalyze the removal of Ub-moieties for their recycling. DUBs serve to counterbalance ubiquitination reactions within the cell, thus dynamically contributing to the regulation of various cellular processes, such as endosomal sorting ^11, 16–18^.

Ub increases proteome complexity, providing an additional surface for protein-protein interactions and functional regulation ^19, 20^. Ub-modified proteins and Ub chains are recognized by several Ub receptors including a family of specialized proteins carrying Ub binding domains (UBDs). To date, over 20 UBD families have been identified and characterized in mammals participating in recognition of Ub monomers or Ub chains on specific substrates ^4, 14, 21^. Together, the Ub network offers a variety of choices to modulate cellular processes including protein membrane trafficking. However, the contribution of the ubiquitinome regarding their impact in intracellular membrane trafficking is not yet fully understood.

To investigate the contribution of most of the members of the ubiquitinome in protein membrane trafficking, we performed High-Content siRNA Screening (HCS) including 1,187 genes of the human “ubiquitinome”. For this screening we used a stable cell line expressing Amyloid Precursor Protein (APP) fused to EGFP. APP was the protein selected as target because it traffics dynamically through the secretory and endocytic pathways containing specific sorting signal motifs ^22–24^ and specific lysine residues targets of ubiquitination ^25–28^. We thus identified the deubiquitinating (DUB) enzyme PSMD14, a subunit of the 19S regulatory particle (RP) of the proteasome, as a crucial player of Golgi-to-Endoplasmic reticulum (ER) retrograde transport. The DUB enzyme PSMD14 has been shown to be specific for K63-Ub chains in cells ^29^. Here, we found that the inhibition of its activity blocks Golgi-to-ER retrograde transport causing the swelling of the Golgi apparatus. Also, we found that PSMD14 inhibition acts as a potent blocker of macroautophagy as a result of its negative impact in Golgi-to-ER retrograde transport. Inhibition of the proteolytic core of the 20S proteasome by MG132 did not recapitulate these effects, indicating that the 19S RP plays a role on macroautophagy by controlling membrane trafficking at the early secretory pathway.

## Results

### High-Content siRNA Screening revealed PSMD14 deubiquitinating enzyme as a novel regulator of protein trafficking

To investigate the contribution of the ubiquitinome in protein membrane trafficking, we performed an automated HCS using a human small interfering RNA (siRNA) “ubiquitinome” library, consisting of 1,187 siRNA duplex pools targeting all known and predicted Ub-genes. This library, used previously in mammalian screening assays ^30, 31^ includes E1 activating enzymes (0.8%), E2 conjugating enzymes (4.7%), E3 Ub ligases (61.2%), UBD-containing proteins (12.8%), DUBs (8.8%), SUMO-specific proteases (SENPs) (0.2%) and others (11.4%) (Fig. 1A). We used this library to identify novel regulators of protein membrane trafficking using an H4 neuroglioma cell line stably expressing APP-EGFP, a cell line previously characterized and used in siRNAs knockdown experiments ^24, 32, 33^. We standardized the basal (background) and maximum fluorescence intensity by analyzing total fluorescence of silenced and non-silenced APP-EGFP, respectively. A reduction in total fluorescence intensity from 100% to 6% was observed with the EGFP siRNA, compared to the Non-Target (NT) siRNA with a Z factor of 0.69 (Fig. 1B). Representative images of basal and maximum levels of total fluorescence intensity in reporter cells are shown in Fig. 1C. Cells were further assessed with a primary siRNA screening consisting of a pool of four-different siRNA duplexes for each of the 1,187 genes selected, each siRNA targeting a distinct region to minimize their off-targets effects ^34^. Fig. 1D shows a summary of the results with all siRNA pools tested and normalized in respect to the expression of the NT siRNA. Among the 1,187 tested genes, we found that the highest total fluorescence increase (4.15-fold increase) corresponded to protein PSMD14 (POH1, also known as Rpn11/MPR1/SS13/CepP1) (Fig. 1D), a subunit of the 19S regulatory particle (RP) of the proteasome, which has DUB activity ^35, 36^. Representative images of reporter cells knock-down (KD) of PSMD14 in comparison to cells transfected with NT siRNA are shown in Fig. 1E. Further, we validated these results analyzing the effect of PSMD14 KD on the levels of full-length endogenous APP in parental H4 cells by western blot. The effect of all four PSMD14 siRNAs in KD cells tested on endogenous APP levels are shown in Fig. 2A (first panel, lanes 3-6) and compared to un-transfected cells (Mock) or cells transfected with NT siRNA (Fig. 2A, first panel, lanes 1 and 2, respectively). Quantification analysis showed a 3.13-fold increase of full-length endogenous APP levels in PSMD14 KD cells, compared to Mock cells (Fig. 2B). Besides, we found that silencing of PSMD14 caused a strong increase in high molecular weight Ub conjugates, consistent with the role of PSMD14 as a proteasomal subunit with DUB activity (Fig. 2A, second panel, lanes 3-6 compared to lanes 1 and 2). Confirmation of the efficient KD of PSMD14 with all four siRNAs tested was demonstrated by western blot (Fig. 2A, third panel, lanes 3-6 compared to lanes 1 and 2), observing a 5.22-fold decrease on PSMD14 levels (Fig. 2C). Moreover, efficient KD of PSMD14 was also determined by quantitative reverse transcription PCR (RT-qPCR) (Fig. 2D). Finally, to evaluate whether the increase of APP protein levels in PSMD14 KD cells could be the result of an up-regulation of APP transcription, we evaluated *APP* mRNA levels by RT-qPCR, observing no significant changes (Fig. 2E). Altogether, these findings indicate that the silencing of PSDM14 caused a robust increase of APP protein levels, a finding that could suggests impairment in its turnover due to trafficking alterations.

**Figure 1.**
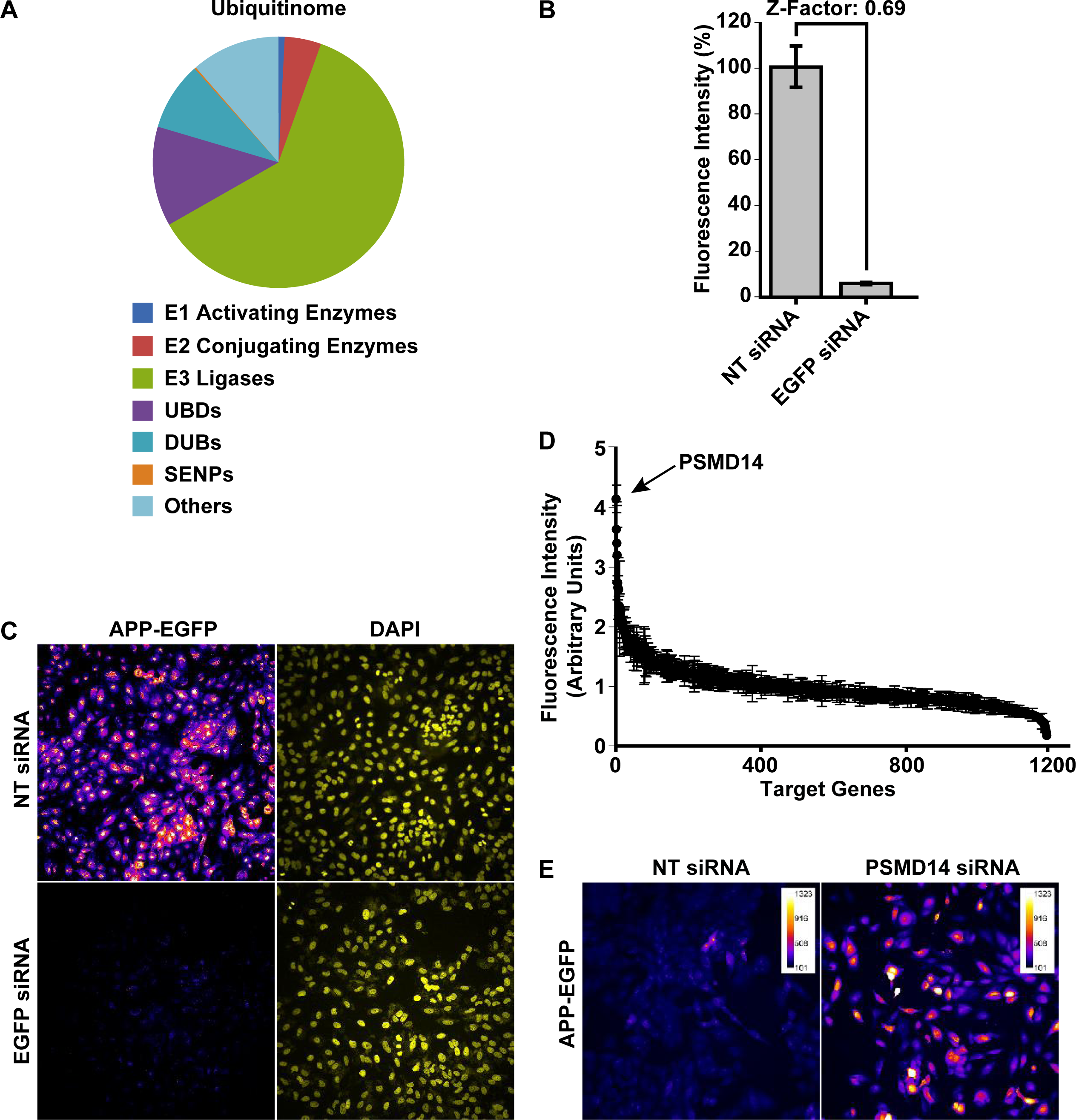
High-Content siRNA Screening assay revealed PSMD14 as a novel regulator of APP levels. **(A)** Graphical distribution of the targets evaluated in the primary High Content siRNA screening using the siRNA “ubiquitinome” library in H4 cells. **(B)** Quantification of the total fluorescence intensity of reporter APP-EGFP cells transfected for 72 h with NT siRNA and EGFP siRNA. Bars represent the mean ± SD with a statistical Z factor = 0.69. **(C)** High content images (20X) captured in reporter APP-EGFP cells transfected for 72 h with NT siRNA and EGFP siRNA. **(D)** Graphical representation of total fluorescence intensity of all 1,187 genes analyzed in primary siRNA screening with the reporter APP-EGFP cells. PSMD14 appears indicated as the top hit. **(E)** High content images (20X) in pseudo color of reporter APP-EGFP cells transfected for 72 h with siRNA SMARTpool targeted against PSMD14 (PSMD14 siRNA) in comparison to cells transfected with NT siRNA. Scale Bar of the images indicates the scale of fluorescence intensity.

**Figure 2.**
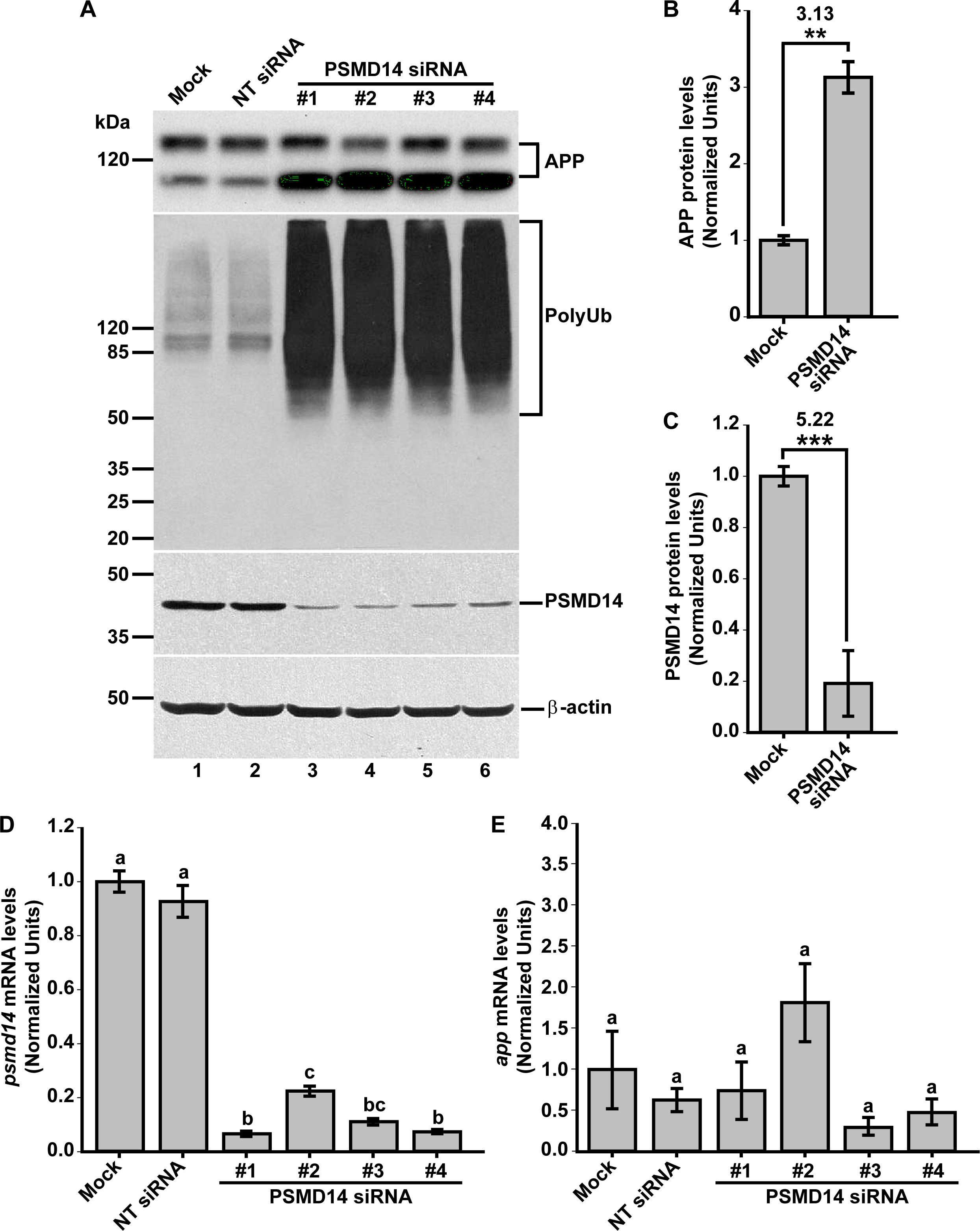
PSMD14 is validated as a regulator of the endogenous APP levels. **(A)** Protein extracts of parental H4 cells either untransfected (Mock), transfected with NT siRNA, or transfected with four different PSMD14 siRNA sequences for 72 h were analyzed by western blot. Polyclonal antibodies to endogenous APP (CT695) and to Ubiquitin, and monoclonal antibodies to PSMD14 (clone D18C7) and to β-actin (clone BA3R), were tested. The position of molecular mass markers are indicated on the left. Densitometric quantification of the levels of endogenous APP protein levels **(B)** and PSMD14 **(C)** in H4 cells transfected with PSMD14 siRNA#1, compared to untransfected cells (Mock). Statistical significance was determined by Student’s t-test. Bars represent the mean ± SD of biological replicates (APP n =5; PSMD14 n=4). **P <0.01 and ***P <0.001. **(D)** mRNA levels of *psmd14* and **(E)** mRNA levels of *app* were measured using RT-qPCR from parental H4 cells transfected for 72 h. All data were normalized for TATA binding protein expression in either untransfected cells (Mock), cells transfected with NT siRNA or cells transfected with four different PSMD14 siRNAs duplexes. Statistical significance was determined by One-Way ANOVA, followed by Tukey’s test. Bars represent the mean ± SD of biological replicates (*psmd14* n=3; *app* n=3). Different letters above the mean bars apply to significant differences between groups P <0.01.

### Acute inhibition of the deubiquitinating enzyme PSMD14 of the 19S RP accumulates APP in a swollen Golgi apparatus

To determine whether PSMD14 is involved in the control of endogenous APP levels, we investigated the effect of incubating cells with Capzimin (CZM), a potent and specific inhibitor of the DUB activity of PSMD14^37^, a subunit of the 19S RP of the proteasome (Fig. 3A). We observed that CZM led to a significant increment in full-length endogenous APP levels in a dose-dependent manner, compared to untreated cells (Fig. 3B and 3C). Interestingly, compared to untreated cells, and in contrast to CZM, treatment with a standard concentration of 10 μM MG132, a reversible inhibitor of the β1, β2 and β5 subunits of the 20S catalytic core of the proteasome^38, 39^ (Fig. 3A), caused a significant decrease in full-length endogenous APP levels (Fig. 3B and 3C). To confirm the inhibition of PSMD14 by CZM, we tested the effect of this inhibitor on the levels of high molecular weight Ub conjugates, comparing it with MG132. In agreement with the effect of PSMD14 KD, we observed that CZM caused a robust increase in high molecular weight Ub conjugates in a dose-dependent manner, finding a maximum effect with 10 μM CZM (Fig. 3D). However, we noticed that 10 μM MG132 caused a more powerful increase in Ub conjugates (Fig. 3D). To confirm the accumulation of Ub conjugates by these treatments, we studied the presence of Ub-inclusion bodies by immunofluorescence analysis. We found that, compared to untreated cells (Fig. 3E, left panel), 10 μM CZM (Fig. 3E, middle panel) caused a similar accumulation of cytoplasmic Ub-inclusion bodies than 10 μM MG132 (Fig. 3E, right panel). Together, our results confirm that acute inhibition of PSMD14 by CZM replicates the phenotype obtained by PSMD14 KD regarding to the impact on the endogenous APP levels. Thus, CZM offers a pharmacological tool to address whether the activity of PSDM14 is required for APP membrane trafficking.

**Figure 3.**
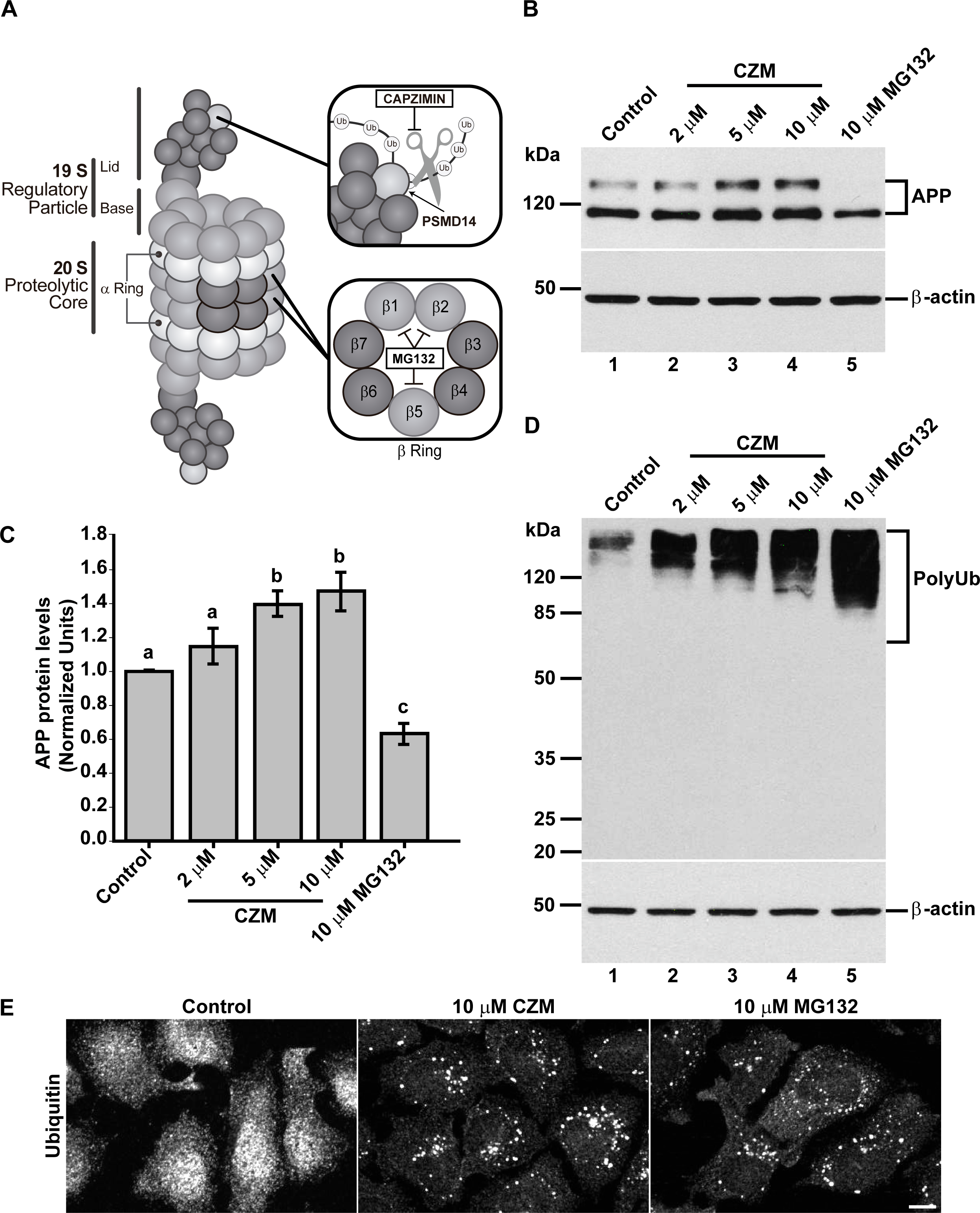
Acute inhibition of PSMD14 by CZM shows a similar phenotype as PSMD14 KD on the levels of APP and high molecular weight Ub conjugates. **(A)** Schematic diagram of the molecular targets of Capzimin and MG132 in the 19S RP and 20S catalytic core of the proteasome, respectively. **(B)** Parental H4 cells were treated either with vehicle (DMSO; Control), or increasing doses of CZM for 4 h, or MG132 for 6 h. Protein extracts were analyzed by western blot with a polyclonal antibody to endogenous APP. Monoclonal antibody to β-actin (clone BA3R) was used as a loading control. The position of molecular mass markers is indicated on the left. **(C)** Densitometric quantification of APP protein levels as shown in (D). Statistical significance was determined by One-Way ANOVA, followed by Tukey’s test. Bars represent the mean ± SD of biological replicates (n=4). Different letters above the mean bars apply to significant differences between groups P <0.05. **(D)** Parental H4 cells were treated as in (B), and the protein extracts were analyzed by western blot with a polyclonal antibody to Ub. Monoclonal antibody to β-actin (clone BA3R) was used as a loading control. The position of molecular mass markers is indicated on the left. **(E)** Immunofluorescence microscopy images of the cellular localization of Ub in parental H4 cells treated with either the vehicle (DMSO; Control), CZM for 4 h or MG132 for 6 h. Cells were fixed, permeabilized and stained with a mouse monoclonal antibody to Ub (clone P4D1) followed by Alexa-488-conjugated donkey anti-mouse IgG. Scale bar, 10 μm. (n=3).

To unveil this possibility, we performed an immunofluorescence analysis of endogenous APP in parental H4 cells. We observed that the treatment with 10 μM CZM resulted in a perinuclear redistribution of APP, which is highly indicative of Golgi apparatus localization (Fig. 4D, compared to 4A). Moreover, similar to the PSMD14 KD, we observed that the CZM treatment caused a significant 2.50-fold increase in APP total fluorescence intensity compared to untreated cells (Fig. 4G). We also observed a significant 1.92-fold increase in the amount of APP in the area positive to the Golgi matrix protein GM130 (GM130), compared to the total area (Fig. 4F, compared to 4C and Fig. 4H). Moreover, we observed that the treatment with CZM caused a significant 1.30-fold increase in the total cell area (Fig. 4I). In addition, we found that CZM caused the swelling of the Golgi apparatus (Fig. 4E compared to 4B), a phenotype that is shown in a zoom in Fig. 4J. To confirm this phenotype, we performed measurements of GM130 images from confocal 3D reconstructions from the Z-stacks of cells treated with CZM (Fig. 4K). We observed a 1.53-fold increase in the Golgi apparatus volume in CZM treated cells, compared to control cells (Fig. 4L). Similar results were observed in quantitative 2D image analysis, observing a significant 2.53-fold increase in the Golgi apparatus area upon CZM treatment (Suppl. Fig. 1). Altogether, our results show that acute inhibition of PSMD14 by CZM accumulates APP in a swollen Golgi apparatus. These findings strongly suggest that CZM impairs the Golgi apparatus due to perturbations of the trafficking through this organelle.

**Figure 4.**
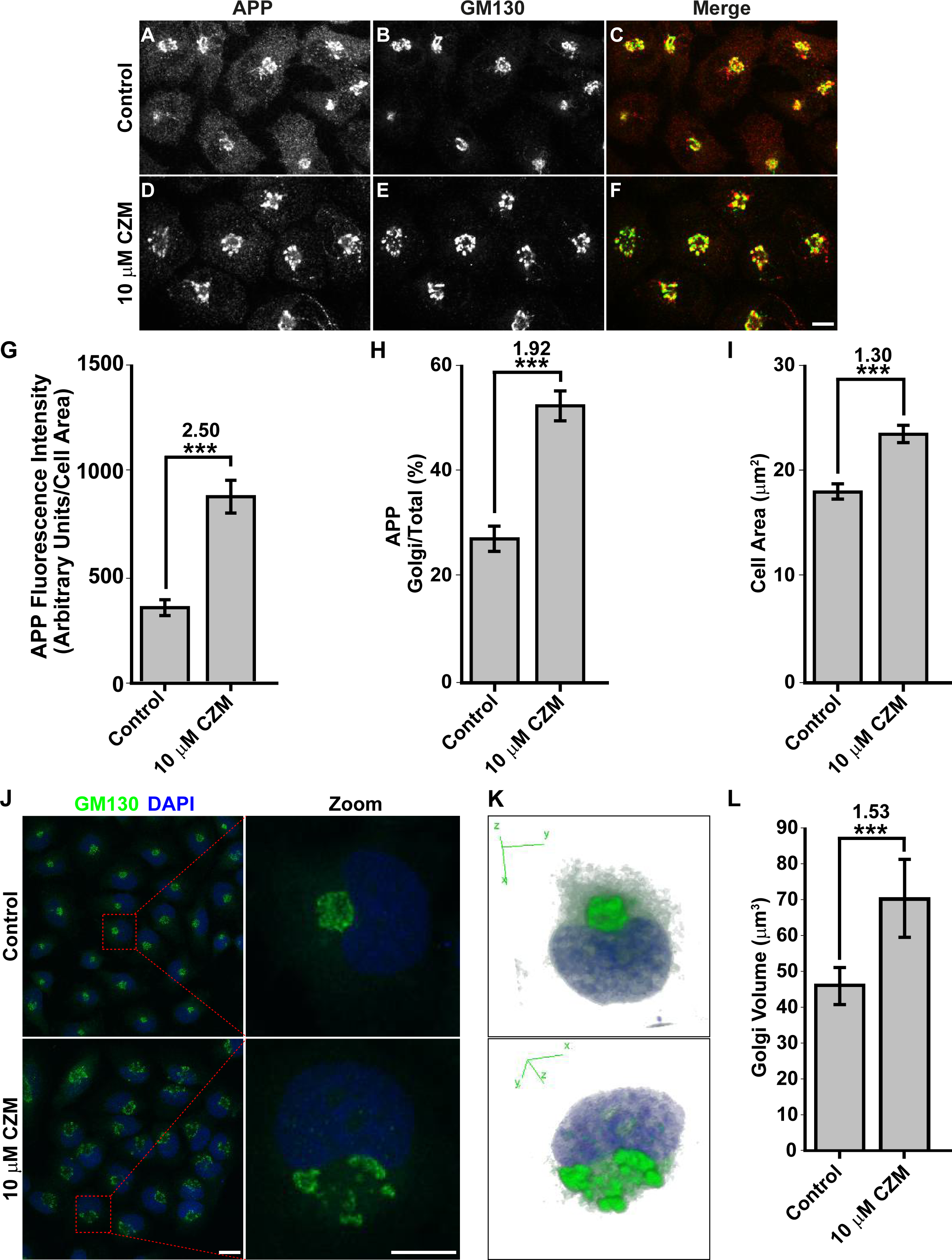
Acute inhibition of PSMD14 by CZM triggers the accumulation of APP in a swollen Golgi apparatus. Immunofluorescence analysis of endogenous APP in H4 parental cells treated either with the vehicle (DMSO; Control) **(A-C)** or CZM **(D-F)** for 4 h. Cells were fixed, permeabilized, and double stained with a rabbit polyclonal antibody to APP (CT695) (A and D) and a mouse monoclonal antibody to GM130 (clone35/GM130) (B and E), followed by Alexa-594-conjugated donkey anti-Rabbit IgG and Alexa-488-conjugated donkey anti-Mouse IgG. Merging of the images generated the third picture (C and F). Scale bar, 10 μm. **(G)** Quantitative analysis of the mean of total fluorescence intensity of APP upon treatment with CZM, in comparison to control cells. The statistical significance was determined by Student’s t-test. Bars represent the mean ± SD of the fluorescent signal per cell area (n=43 cells). ***P <0.001. **(H)** Quantitative analysis of the fraction of APP colocalizing with GM130 under CZM treatment and compared to control cells. Statistical significance was determined by Student’s t-test. Bars represent the mean ± SD of the fluorescent signal per cell area (n=43 cells). ***P <0.001. **(I)** Quantitative analysis of the cell area. Statistical significance was determined by Student’s t-test. Bars represent the mean ± SD of the cell area (n=43 cells) **P <0.001. **(J)** Immunofluorescence microscopy analysis of GM130 in parental H4 cells treated either with the vehicle (DMSO; Control) or CZM for 4 h. Cells were fixed, permeabilized and stained with mouse monoclonal antibody to GM130 (clone 35/GM130) followed by Alexa-488-conjugated donkey anti-mouse IgG, and nuclei were stained with DAPI. Scale bar, 10 µm. **(K)** 3D reconstructions of the Golgi apparatus using GM130 as Golgi marker were generated from Z-stacks (250 nm). **(L)** Golgi Volume was measured from 3D reconstructions as shown in (K). Statistical significance was determined by Student’s t-test. Bars represent the means ± SEM (n=20 cells). *** P <0.001.

### Acute inhibition of the deubiquitinating enzyme PSMD14 perturbs Golgi-to-ER retrograde transport

The Golgi apparatus is a highly dynamic organelle that requires fine regulation of trafficking pathways in order to maintain its size, shape and composition. In particular, it has been shown that Golgi-to-ER retrograde transport plays a crucial role in the maintenance of Golgi morphology. Indeed, inhibition of this specific trafficking pathway results in a significant swelling of this organelle ^40–45^. To investigate whether the swelling of the Golgi by the inhibition of PSMD14 with CZM could be the result of the inhibition of the Golgi-to-ER retrograde transport, we first validated our findings in HeLa cells. We measured Golgi apparatus volume in 3D reconstructions from the Z-stacks of untreated HeLa cells or treated with CZM and MG132 by using Giantin as Golgi apparatus reporter. Similar to our findings in H4 cells, we found that the inhibition of PSMD14 DUB activity by CZM causes a 2.14-fold increase in Golgi apparatus volume (Fig. 5A, middle panel compared to the left panel and 5B). Interestingly, in contrast to CZM, we observed that MG132 caused no effect on Golgi apparatus volume (Fig. 5A, right panel compared to the left panel and 5B), strongly suggesting that Golgi homeostasis is regulated by the PSMD14 DUB activity. Then, we investigated whether this phenotype was the result of Golgi-to-ER retrograde transport inhibition. We assessed the distribution of stably overexpressed KDEL (Lys-Asp-Glu-Leu) Receptor 1 (KDELR1) fused to GFP (KDELR1-GFP) in HeLa cells. As it was previously reported ^44, 46^, overexpressed KDELR1-GFP is mostly distributed to the ER due to its efficient transport from the Golgi apparatus to the ER (Fig. 5C, left panel and 5D). Thus, impairment of Golgi-to-ER retrograde transport causes the accumulation of KDELR1-GFP in the Golgi apparatus working as an assay to identify novel regulators of this pathway ^44^. Interestingly, we found that CZM caused a rapid time-dependent accumulation of KDELR1-GFP at the Golgi apparatus, observing a maximal effect after 90 min of treatment (Suppl. Fig. 2). At this time, we observed a significant 1.50-fold increase in the amount of KDELR1-GFP within the Giantin-positive Golgi apparatus, compared to the total area (Fig. 5C, middle panel compared to the left panel, and 5D). As before, we tested the effect of MG132, observing no effect on the distribution of KDELR1-GFP (Fig 5C, right panel compared to left panel, and 5D), which strongly supports that MG132 has no effect on Golgi-to-ER retrograde transport. Next, to confirm blockage of Golgi-to-ER retrograde transport by the acute inhibition of the PSMD14, we tracked the redistribution of the transiently overexpressed thermo-sensitive KDELR1-VSVG-YFP chimera (vesicular stomatitis virus G protein fused to KDEL receptor 1 and YFP) in HeLa cells upon treatment with CZM by live cell imaging at different temperatures. Briefly, at a permissive temperature of 32°C, KDELR1-VSVG-YFP cycles between the Golgi apparatus and the ER, showing distribution mainly at the Golgi apparatus. Upon shifting to a restrictive temperature of 40°C, KDELR1-VSVG-YFP is progressively accumulated at the ER due to its specific retention in this compartment at this temperature. In addition, because the Golgi-to-ER retrograde transport is not affected at 40°C, the rapid decay of the fluorescence of KDELR1-VSVG-YFP fluorescence at the Golgi apparatus, a measurement that is used to evaluate inhibition of Golgi-to-ER retrograde transport pathway ^47^. We found that in control cells nearly 50% of the fluorescence of KDELR1-VSVG-YFP decayed after 15 min of shifting the temperature to 40°C (Fig. 5E, upper panels and 5F), indicating that retrograde transport is working normally ^47^. In contrast, when cells were treated with CZM, the decay of the fluorescence of KDELR1-VSVG-YFP was not apparent, confirming that acute inhibition of the PSMD14 blocked Golgi-to-ER retrograde transport (Fig. 5E, lower panels and 5F). In contrast, MG132 treated cells showed similar results as controls (data not shown), confirming that the 20S proteasome has no effect on retrograde transport. Importantly, we found that CZM is able to block retrograde transport for short times having no effect on the catalytic activity of the 20S proteasome (Suppl. Fig. 3). In contrast, a rapid and robust inhibition on the catalytic activity of the 20S proteasome is observed with MG132 (Suppl. Fig. 3). These findings strongly indicate that acute inhibition of PSMD14 DUB activity by CZM acts as a powerful blocker of Golgi-to-ER retrograde transport, explaining the swelling of the Golgi apparatus and the accumulation of protein cargoes such as APP at this location.

**Figure 5.**
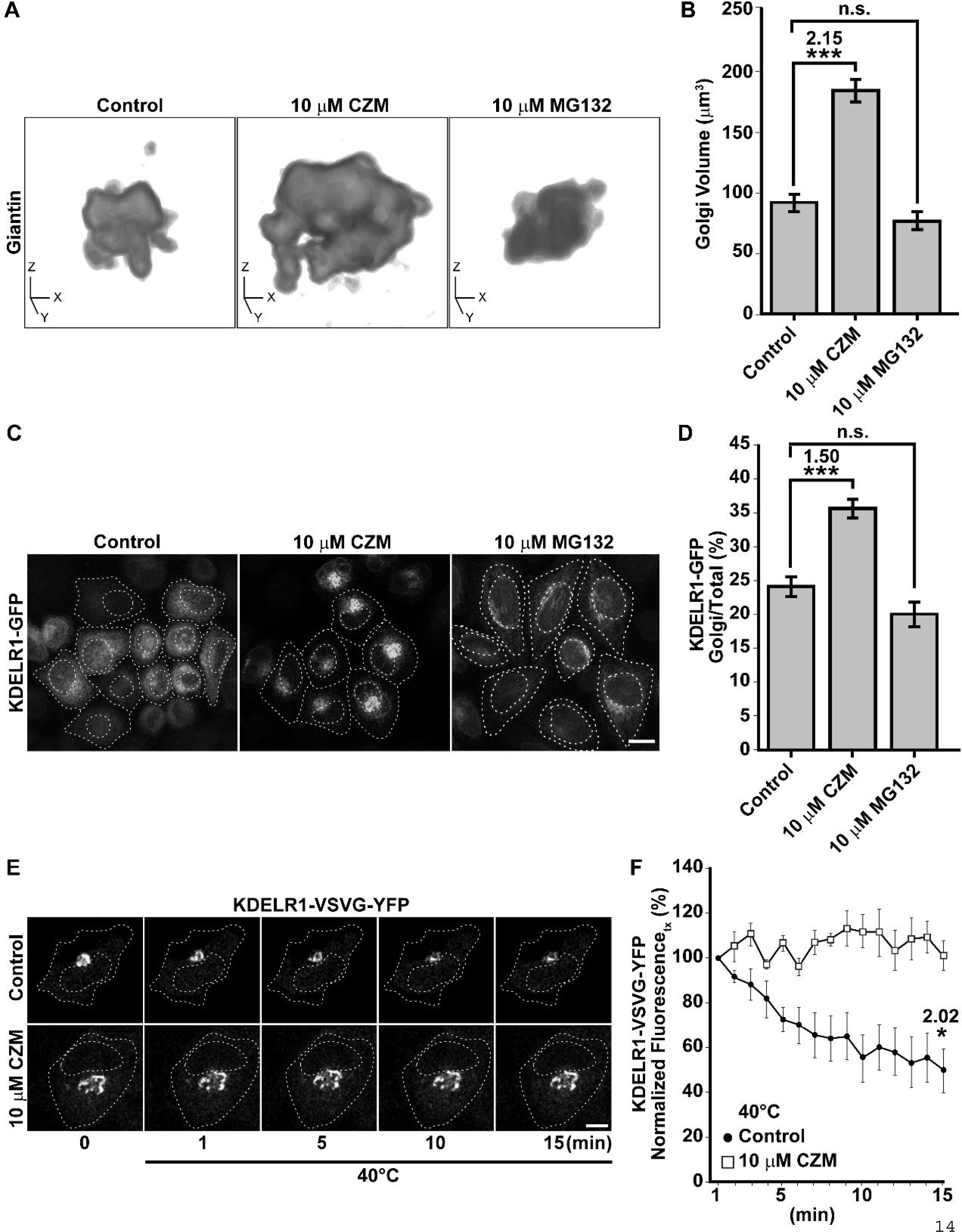
The PSMD14 DUB inhibitor CZM impairs Golgi-to-ER retrograde transport. **(A)** 3D reconstructions of the Golgi apparatus using Giantin as Golgi marker were generated from Z-stacks (250 nm) obtained from HeLa cells stably expressing KDELR1-GFP treated for 90 min either with vehicle (DMSO; Control), CZM or MG132. **(B)** Golgi volume was measured from 3D reconstructions as shown in (A). Statistical significance was determined by Student’s t-test. Bars represent the means ± SEM (n=30 cells). *** P <0.001. **(C)** HeLa cells stably expressing KDELR1-GFP were treated for 90 min either with vehicle (DMSO; Control), CZM or MG132. Cells were fixed and representative confocal images were acquired. **(D)** Measurement of Giantin and total KDELR1-GFP total fluorescent intensity. Statistical significance was determined by Student’s t-test. Bars represent the means ± SEM (n=34 cells). *** P <0.001. **(E)** H4 cells were transiently transfected to express the thermo-sensitive retrograde transport reporter KDELR-VSVG-YFP. Cells were kept at 32°C to allow KDELR-VSVG-YFP localization at the Golgi. Cells were then shifted to 40°C (restrictive temperature) and images acquired at 1 min interval for 15 min. **(F)** Quantitative image analysis was performed to measure the integrated fluorescence of KDELR-VSVG-YFP at the Golgi at 1 min interval for 15 min. Statistical significance was determined by Student’s t-test. Bars represent the mean ± SEM (n=3 cells). *p<0.05

### Inhibition of Golgi-to-ER retrograde transport by CZM has a negative impact on macroautophagy

Several lines of evidence have shown that Golgi-to-ER retrograde transport plays a relevant role in autophagosome biogenesis at the level of the ER ^48–51^. In addition, it has been shown that PSMD14 participates in the activation of the aggresome clearance by cleaving K63 Ub chains of aggregate proteins^52, 53^. Thus, we investigated the effect of acute inhibition of PSMD14 DUB activity on the number of autophagosomes by testing the classical marker microtubule-associated protein 1 light chain 3B (LC3B) ^54^. Therefore, parental H4 cells were treated under nutrient starvation with Earle’s balanced salt solution (EBSS), an established culture medium used for activation of autophagosomal formation ^55^. As expected, we found that starvation strongly increased the number of autophagosomes (Fig. 6B), compared to cells treated under normal nutrient conditions (Fig. 6A). In contrast, we observed that treatment with CZM abolished the appearance of autophagosomes upon EBSS treatment (Fig. 6C compared to 6B). We also tested the effect of CZM in cells under normal nutrients conditions, but in the absence or presence of Torin-1, a potent and selective inhibitor of mammalian target of rapamycin complexes (mTORC1/2), a trigger of autophagosomal formation^56^. Similar to the results with EBSS, we observed that CZM prevented the appearance of autophagosomes promoted by Torin-1 (Fig. 6F compared to 6E). In agreement with these findings, we observed that CZM alone did not increase the number of autophagosomes (Fig. 6D compared to 6A), in contrast to the effect of MG132 (Suppl. Fig. 4). To confirm these results, we quantified the number of LC3-positive structures under all conditions tested, observing a significant decrease in the number of autophagosomes when cells, treated with EBSS solution or Torin-1, where also treated with CZM (Fig. 6G). In addition, we validated these results biochemically performing western blot analysis of endogenous LC3B (Fig. 6H). We found that the levels of LC3B-II were strongly increased with EBSS or Torin-1 treatment (Fig. 6H, lanes 2 and 4). In contrast, when cells were treated with EBSS or Torin-1 in the presence of CZM, LC3B-II levels did not change (Fig. 6H, lanes 3 and 5) compared to control cells (Fig. 6H, lane 1), results that were quantified and depicted in Fig. 6J. Interestingly, this biochemical analysis also showed that CZM increased LC3B-I levels under all conditions tested (Fig. 6H, lanes 3, 5 and 6, and Fig. 6I) compared to control cells (Fig. 6H, lane 1), suggesting that the reduction of the autophagosomal structures caused by CZM was not due to a reduction in LC3B total levels. Altogether, these results confirm that acute inhibition of PSMD14 DUB activity acts as a potent blocker of autophagosome biogenesis induced by EBSS or Torin-1. In addition, these findings suggest that blockage of autophagosomal biogenesis could be a consequence of Golgi-to-ER retrograde transport inhibition.

**Figure 6.**
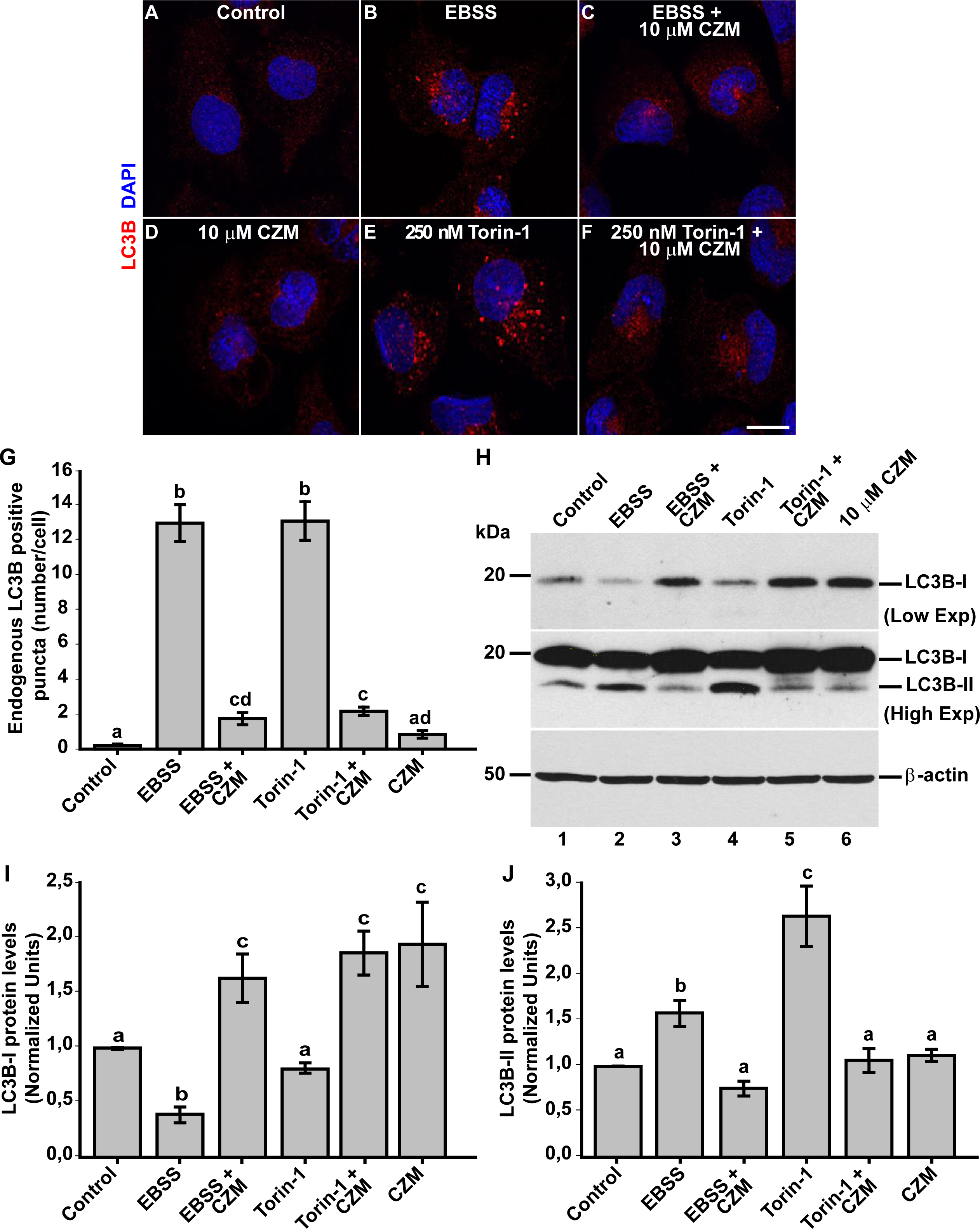
Inhibition of autophagosome formation by CZM. Immunofluorescence microscopy analysis of the subcellular localization of endogenous LC3B in parental H4 cells treated with either the vehicle (DMSO; Control) **(A)**, EBSS for 4 h **(B),** CZM for 6 h **(D)** or Torin-1 for 4 h **(E)**. EBSS **(C)** and Torin-1 **(F)** were tested using a 2-h pretreatment with CZM followed by the treatment with EBSS or Torin-1 for 4 h in the presence of CZM. Cells were fixed, permeabilized and stained with a rabbit polyclonal antibody to LC3B followed by Alexa-594-conjugated donkey anti-Rabbit IgG, and nuclei were stained with DAPI. Scale bar 10 µm. **(G)** Quantification of the puncta positive to LC3B. Statistical significance was determined by One-Way ANOVA, followed by Tukey’s test. Bars represent the mean ± SEM (n=50 cells). Different letters above the mean bars indicate the significant differences between groups P <0.05. **(H)** Protein extracts from parental H4 cells treated as in (A-F) were analyzed by western blot with a rabbit polyclonal antibody to LC3B. Monoclonal antibody to β-actin (clone BA3R) was used as a loading control. The position of molecular mass markers is indicated on the left. **(I)** Densitometric quantification of LC3B-I levels and **(J)** LC3B-II levels. Statistical significance was determined by One-Way ANOVA, followed by Tukey’s test. Bars represent the mean ± SEM of biological replicates (LC3B-I n=3; LC3B-II n=3). Different letters above the mean bars indicate the significant differences between groups P <0.05.

### Inhibition of Golgi-to-ER retrograde transport by CZM accumulates RAB1A and ATG9A at the Golgi apparatus

To evaluate this hypothesis, we tested the effect of acute inhibition of the PSMD14 DUB activity on the distribution of proteins implicated in the initial steps of autophagosome formation, which traffics in early compartments of the secretory pathway. We first tested RAB1A, a small GTPase with an essential role in the initiation of autophagy, facilitating the recruitment of the unc-51-like kinase 1 (ULK1) complex to subdomains of the ER, a crucial early step during autophagosome formation ^57, 58^. We found that CZM treatment caused a redistribution of RAB1A to the perinuclear zone (Fig. 7A). Measuring these images confirmed a significant increase of RAB1A in this area (Fig. 7B), a result that was accompanied by a decrease of RAB1A in the cell periphery defined as radial zone (Fig. 7B). Importantly, we found that RAB1B was not affected by the same condition (data not shown). Moreover, we observed that RAB1A was distributed in the swollen Golgi apparatus, similar to GM130, upon CZM treatment (Suppl. Fig. 5). With the same aim, we studied the distribution of ATG9A, an essential transmembrane protein involved in macroautophagy, which plays a crucial role in the early steps of autophagosome formation ^59, 60^. ATG9A-containing vesicles are formed from the Golgi apparatus translocating to the ER to form the initiation site at the ER for autophagosome formation ^61^. Similar to RAB1A, we found that acute inhibition of PSMD14 caused a redistribution of ATG9A to the perinuclear zone, together with a decrease in its distribution in the radial zone (Fig. 7C). The quantification analysis of these images is shown in Fig. 7C. Moreover, and similar to RAB1A, we observed that ATG9A is distributed to the swollen GM130-Golgi apparatus area upon CZM treatment (Suppl. Fig. 6). These results strongly indicate that blockage of Golgi-to-ER retrograde transport by acute inhibition of PSMD14 DUB activity causes the retention at the Golgi apparatus of key proteins implicated in early steps of autophagosome formation. The PSDM14-dependent inhibition of autophagy explains the accumulation of APP at the Golgi apparatus since macroautophagy has been recently demonstrated as a positive regulator of protein secretion from the Golgi apparatus^62^. Collectively, these results show the strong interplay between membrane transport and autophagy through a novel mechanism involving the proteasome complex through the deubiquitinating activity of PSMD14.

**Figure 7.**
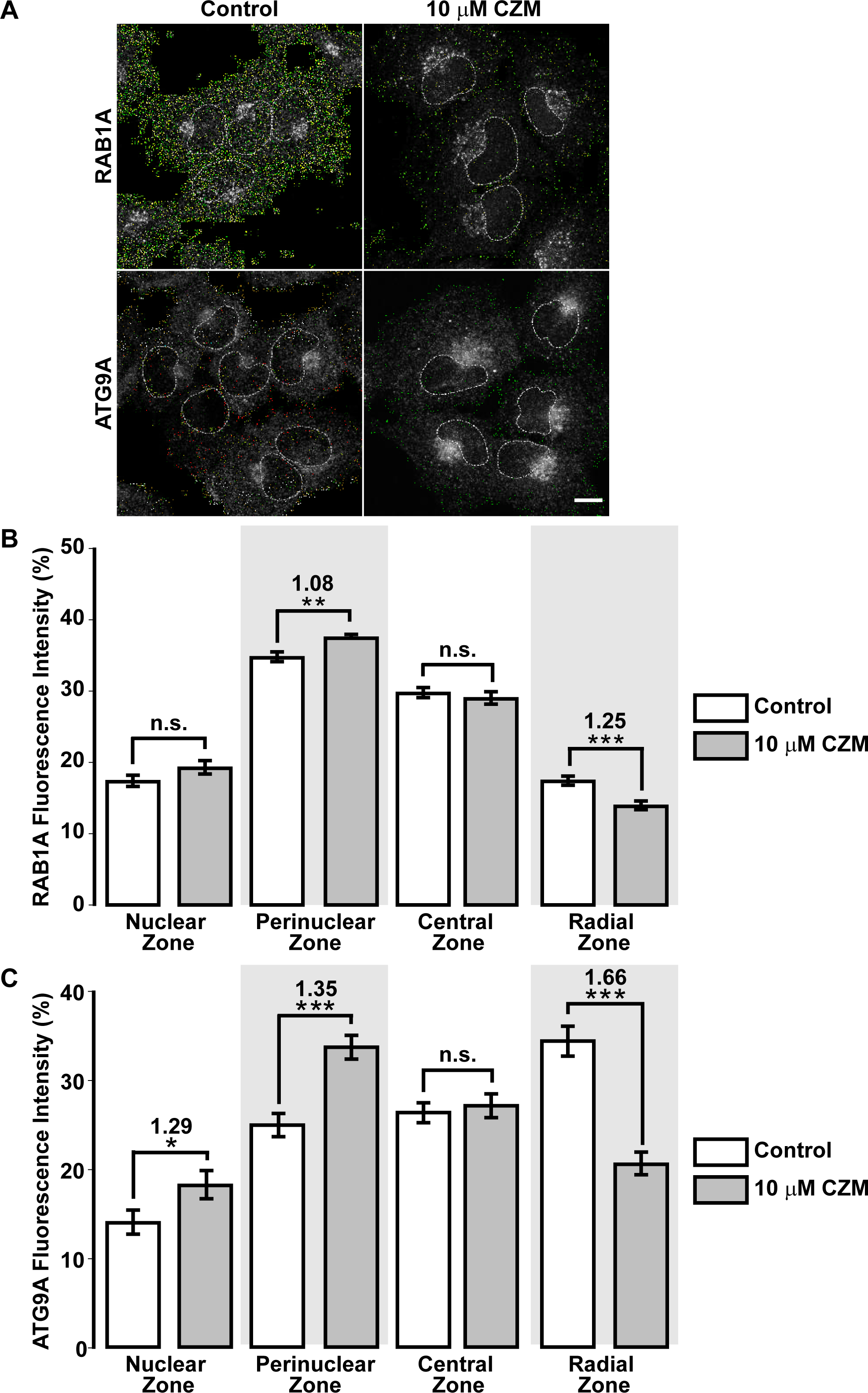
Redistribution of RAB1A and ATG9A to the Golgi apparatus with CZM. **(A)** Immunofluorescence analysis of endogenous RAB1A and ATG9A in H4 parental cells treated for 4 h either with the vehicle (DMSO; Control) (left panel) or CZM (right panel). Cells were fixed, permeabilized, and stained with a rabbit monoclonal antibody to RAB1A (clone D3X9S) (upper panel) and a rabbit monoclonal antibody to ATG9A (clone EPR2450(2)) (lower panel), followed by Alexa-594-conjugated donkey anti-Rabbit IgG. Scale Bar, 10 μm. (B) Quantitative analysis of the fluorescence intensity of RAB1A upon treatment with CZM, in comparison to control cells. Statistical significance was determined by Student’s t-test. Bars represent the mean ± SEM of the fluorescent signal per cell area (n=227 cells). **P <0.01; ***P < 0.001; n.s., not significant. (C) Quantitative analysis of the fluorescence intensity of ATG9A upon treatment with CZM, in comparison to control cells. Statistical significance was determined by Student’s t-test. Bars represent the mean ± SEM of the fluorescent signal per cell area (n=95 cells). *P< 0.05; ***P < 0.001; n.s., not significant.

## Discussion

We report here that PSMD14 DUB activity, a subunit of the 19S RP of the proteasome, functions as a novel regulator of autophagosome formation. To our knowledge, this is the first report demonstrating that impairment of the proteasome can have a negative impact on the initiation of autophagy. Several reports have shown that inhibition of the 20S catalytic core of the proteasome by blockers of the β-subunits trigger the enhancement of the biogenesis of LC3B-positive autophagosomes ^63–68^. Contrary to these findings, we found that the blockage of the PSMD14 DUB activity perturbs the biogenesis of LC3B-positive autophagosomes. Because the inhibition of PSMD14 DUB activity but not the blockage of the 20S catalytic core blocked Golgi-to-ER retrograde transport, a pathway implicated in the initiation of autophagosomes ^48–51^, we postulated that PSMD14 DUB activity controls autophagy by a process independent of bulk proteasomal degradation but dependent on K63-Ub chains. Indeed, K63-Ub chains has been directly involved in the recycling of transmembrane cargoes into the endosomal pathway in *C. elegans*^69^. In addition, K63-Ub chains regulate positively autophagy by mTORC inactivation through K63-linked polyubiquitination of RagA^70^, and by promoting K63-chain autoubiquitination of the E3 ligase TRIM13 that increases the interaction with the autophagy receptor p62 promoting initiation of ER-phagy^71^. Likewise, the deconjugation of K63-Ub chains has biological relevance. Free unanchored K63-Ub chains release in a PSMD14 dependent manner are crucial to coordinate the elimination of protein aggregates by autophagy^52^. Free unanchored K63-Ub chains bind and activate the deacetylase HDAC6^52, 53^, which contributes to relocate the aggregates to the aggresome for final autophagic clearence^72–83^. Here, we unveil the first insight into the regulation of Golgi-to-ER retrograde transport by K63-Ub chains by the functional role of the deubiquitinating enzyme PSMD14. We hypothesize that the function of key cytosolic proteins implicated in Golgi-to-ER retrograde transport are under the control of K63-Ub chains, confirming the crucial regulatory role of deubiquitination in membrane protein trafficking ^73, 74^.

In this regard, key regulatory proteins involved in Golgi-to-ER transport are regulated by the state of ubiquitination/deubiquitination. In yeast, deletion of the DUB Ube3p and its co-factor Bre5p accumulates ubiquitinated β’-COP facilitating its rapid degradation by the proteasome ^73, 75^. Because β’-COP is a subunit of the COP-I coatomer complex, a key machinery implicated in Golgi-to-ER retrograde transport, reduction in β’-COP levels perturbs this trafficking pathway ^73^. Two other proteins that participates in this retrograde trafficking pathway, such as PKA and UVRAG ^44, 76^, have been shown to be regulated by ubiquitination. Blockers of PKA signaling cause inhibition of this trafficking pathway, a process that is accompanied by the swelling of the Golgi apparatus ^44, 45^. It has been shown that the catalytic PKA subunit (PKAc) is ubiquitinated by the CHIP E3 ligase, resulting in proteasomal degradation of PKAc and signaling shutdown ^77^. UVRAG mediates the interaction of β’-COP with ER tethers and COP-I coatomer for efficient fusion of retrograde vesicles to the ER, a crucial step during Golgi-to-ER retrograde transport ^49^. UVRAG is ubiquitinated by SMURF1 ^78^ and its silencing causes the swelling of the Golgi apparatus and the inhibition of the Golgi-to-ER retrograde transport ^49^. Whether these proteins are regulated by K63-Ub chains is unknown and should be further addressed.

In addition, we propose that Golgi-to ER retrograde transport inhibition by PSMD14 dysfunction might result on the accumulation of structural and/or autophagy regulatory elements. In fact, the inhibition of PSMD14 caused the accumulation of ATG9A at the Golgi. ATG9A is a multispanning membrane protein essential for autophagy ^59, 60^. ATG9A is actively transported through Golgi-to-ER retrograde transport ^61^ and participates as a key player in the biogenesis of autophagosomes ^59, 79^. Moreover, along with the accumulation of ATG9A at the Golgi, we found increased levels of RAB1A at the Golgi apparatus. RAB1A is an essential small GTPase that participates in the recruitment of the ULK1 complex to subdomains of the ER for autophagy initiation ^57, 58^. Interestingly, and similar to PSMD14 inhibition, ATG9A deficiency causes an increase in LC3B-I levels, accompanied by a reduction in the number of autophagosome structures ^59, 79, 80^. Together, our results strongly support that PSMD14 inhibition perturbs autophagosome biogenesis due to the sequestration of key proteins of this process at the Golgi apparatus. In addition, it supports the hypothesis of a closed intersection between Golgi-to-ER retrograde and autophagy pathway ^49, 76^ whereas PSMD14 DUB activity emerged as a new regulatory element of this intersection. The PSDM14-dependent inhibition of macroautophagy might affect protein secretion from the Golgi apparatus since autophagy has been recently demonstrated as a positive regulator of this process^62^ as observed by the strong accumulation of APP at the Golgi apparatus. Collectively, the results demonstrate the strong functional interplay between membrane transport and macroautophagy mediated by a novel mechanism involving the proteasome complex through the deubiquitinating activity of PSMD14 (Model on Fig. 8). We propose that inhibition of PSDM14 DUB activity by CZM is a new strategy to cause inhibition of the Golgi-to-ER retrograde pathway. Thus, CZM should now be considered a new pharmacological tool to study the impact of Golgi-to-ER retrograde transport inhibition in other cell biology processes, such as autophagy. Moreover, because CZM was discovered as a new drug for cancer treatment ^37^, it would be now interesting to investigate whether part of its anti-cancer effects could be the result of inhibiting Golgi-to-ER retrograde pathway. Interestingly, recent studies have shown that PSMD14 is upregulated (mRNA and protein levels) in different tumoral cell types ^81, 82^. Whether cancer cells are more dependent of Golgi-to-ER retrograde pathway than normal cells, as it happens regarding the mechanisms of protein quality control ^83^ is still unclear. CZM could offer an interesting tool to evaluate this hypothesis positioning PSMD14 as a promising target for therapeutic intervention.

**Figure 8.**
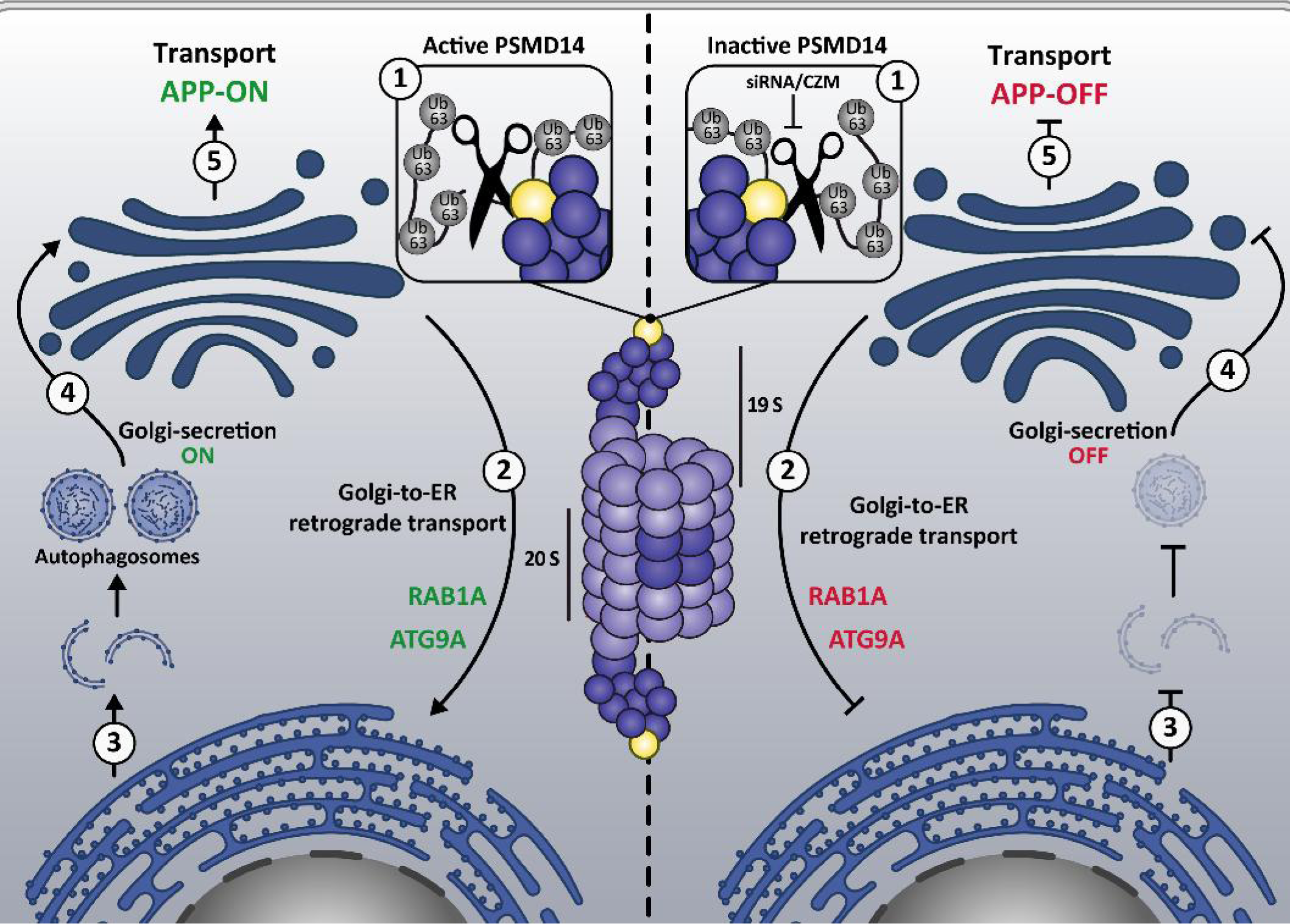
Model of mechanism underlying regulation of protein membrane trafficking and macroautophagy by the proteasome 19S RP PSMD14 DUB activity. The model depicts the closed interplay between membrane transport and macroautophagy by a novel mechanism involving the proteasome complex through the deubiquitinating activity of PSMD14. We propose that active PSMD14 and their K63-Ub chains (1, left panel) positively regulate Golgi-to-ER retrograde transport (2, left panel), pathway implicated in the retrieval of key proteins for autophagosome biogenesis and macroautophagy (3, left panel). Reduction of free K63-Ub chains by inactive PSM14 (1, right panel) results on the blockage of Golgi-to-ER retrograde transport (2, right panel) causing the accumulation of ATG9A and RAB1A at the Golgi apparatus. Thus, blockage of Golgi-to-ER retrograde transport inhibits the biogenesis of autophagosomes and macroautophagy (3, right panel). Macroautophagy has been recently demonstrated to act as a potent positive regulator of protein transport from the Golgi apparatus to the cell surface (Golgi-secretion ON; 4, left panel). Thus, inhibition of macroautophagy upon inactive PSMD14 (siRNA/CZM) blocks protein transport from the Golgi apparatus to the cell surface (Golgi-secretion OFF) explaining the effect on APP transport (5 left and 5 right).

## Materials and Methods

### Chemical Reagents

Earle’s balanced salt solution (EBSS) and the cocktail of protease inhibitors were purchased from Sigma-Aldrich (St. Louis, MO, USA). MG132 was purchased from Millipore (Burlington, MA, USA). Torin-1 was purchased from Tocris Bioscience (Bristol, UK). Dr. Raymond Deshaies from California Institute of Technology (Caltech), CA, USA and Dr. Yuyong Ma from University of California, CA, USA kindly donated Capzimin (CZM).

### Antibodies

We used the following monoclonal antibodies: mouse anti-ubiquitin clone P4D1 (Cytoskeleton, Inc, Denver, Co, USA), mouse anti-β-actin clone BA3R (Thermo Fisher Scientific, Waltham, MA, USA), rabbit anti-RAB1A clone D3X9S, mouse anti-GM130 clone 35/GM130, rabbit anti-PSMD14 clone D18C7 (Cell Signaling Technology, Danvers, MA, USA) and rabbit anti-ATG9A clone EPR2450(2) (Abcam, Cambridge, UK). We used the following polyclonal antibodies: rabbit anti-ubiquitin (cat: Z0458, Dako, Carpintería, CA, USA), rabbit anti-Giantin (cat: AB24586, Abcam, Cambridge, UK), rabbit anti-LC3 (cat: 2775S, Cell Signaling Technology, Danvers, MA, USA), rabbit anti-APP CT695 (cat: 51-2700, Thermo Fisher Scientific). Horseradish Peroxidase-conjugated secondary antibodies were purchased from Jackson ImmunoResearch Laboratories (West Grove, PA, USA), and DAPI probe, Alexa and Dylight fluorophore-conjugated secondary antibodies were purchased from Thermo Fisher Scientific.

### Cell Culture

H4 ATCC ® HTB-148™ Homo sapiens brain neuroglioma cells, referred here as H4 human neuroglioma cells and HeLa ATCC ® CCL-2™ cells were obtained from the American Type Culture Collection (Manassas, VA, USA). Cell lines were cultured in Dulbecco’s modified Eagle’s medium (DMEM; Thermo Fisher Scientific) supplemented with 10% (vol/vol) heat-inactivated fetal bovine serum (FBS; Thermo Fisher Scientific), and penicillin/streptomycin (Thermo Fisher Scientific), in a 5% CO_2_ atmosphere at 37°C. The generation of the H4 stable cell line expressing hemagglutinin-tagged APP_695_-F/P-D/A-EGFP (APP-EGFP) and the HeLa stable cell line expressing KDELR1-GFP were previously reported ^32, 44, 84^. Stably transfected cells were maintained in culture medium supplemented with 100 μg/ml G418. Cells were grown to sub confluence and then treated with drugs or transfected with siRNAs for further western blot and immunofluorescence analyses. Nutrient starvation assays were performed in the presence of EBSS. Assays to detect *Mycoplasm* were performed periodically.

### High Content siRNA Transfection and Imaging

The primary siRNA screen for levels of APP-EGFP were performed in duplicates with our custom-assembled “Ubiquitinome” siRNA library, which consists of 1,187 SMARTpools siRNAs targeting all known and assumed components of the ubiquitin and ubiquitin-like systems in 96-well format, previously reported ^30, 31^. The H4 cells stably expressing APP-EGFP were reverse transfected ^30, 31^ in µClear bottom 96-well plates (Greiner Bio-One, Kremsmünster, Austria) with ON-TARGETplus SMARTpools siRNAs (GE Dharmacon, Lafayette, CO, USA). Additionally, ON-TARGETplus Non-targeting pool (NT siRNA) was used together with untransfected cells (Mock) as neutral control; siRNA against GFP (GFP siRNA) was used as positive control for APP downregulation. Briefly, 10 µl siRNA (200 nM) were stamped from 96-well plates of the library on to µClear bottom 96 well plates. Thereafter, 10 µl of Opti-MEM I Reduced Serum Medium (Thermo Fisher Scientific) containing Lipofectamine RNAiMAX transfection reagent (Thermo Fisher Scientific) was added to the siRNA (dilution 1:50). Plates were shaken at 900 rpm for 1 min followed by incubation for 20 min at room temperature. During this incubation cells were resuspended in DMEM supplemented with 10% (vol/vol) FBS and 1x Normocin (InvivoGen, San Diego, CA, USA) and 6,000 cells were dispensed onto each well loaded with medium for a final siRNA concentration of 20 nM. Plates were incubated for 72 h and further prepared for high content measurement.

After transfection, cells were stained using 100 μl of reagent per step, dispensed by the automated reagent dispenser XPP-721 (fluidX, Manchester, UK), according to the following protocol: two washes in phosphate buffered saline (PBS); fixation in 3.7% (vol/vol) paraformaldehyde for 10 min; one wash in PBS; incubation with 0.2% (vol/vol) Triton X-100 in PBS for 10 min; two washes in PBS; incubation with 0.1 µg/ml DAPI for 5 min; two washes with PBS. Finally, 100 µl of PBS were left in each well. Images were acquired and analyzed using the automated microscope IN Cell 2000 Analyzer (GE Healthcare, Little Chalfont, UK). Images of six randomized fields per well having ∼ 600 cells each were acquired, and further analyzed the total fluorescence intensity in n > 2,000 cells per condition. A secondary siRNA screening was performed in triplicate targeting the 35 most responsive hits, using each single siRNA duplex derived from the SMARTpools used in the primary siRNA screening.

### siRNA Transfection for the siRNA Screening Validation Stage

Four single siRNA sequences targeting human PSMD14 (Accession number: NM_005805), derived from the ON-TARGETplus SMARTpool used in the siRNA Screening (Suppl. Fig. 7) were purchased from GE Dharmacon (Lafayette, CO, USA). siRNA transfections were carried out in 60 mm tissue culture plates using the Lipofectamine RNAiMax transfection reagent (Thermo Fisher Scientific) according to the manufacturer’s protocol, and after 72 h cells were collected for further analysis.

### RNA isolation and RT-qPCR Analysis

Total RNA extraction from H4 cells was carried out using the E.Z.N.A.® Total RNA Kit I (Omega Biotek, Norcross, GA, USA), and either purity (260nm/280nm ratio and 260nm/230nm ratio) and quantity (260nm absorbance) were determined by spectrophotometry using NanoVue Spectrophotometer (GE Healthcare). The cDNA synthesis was performed from 2.5 μg of total RNA and oligo-dT and MMLV reverse transcriptase (Promega, Madison, WI, USA) according to supplier instructions. Specific primer pairs for *tbp* (NM_003194), *psmd14* (NM_005805) and *app* (NM_000484) human genes were designed for quantitative reverse transcription PCR on cDNA template (RT-qPCR) (Suppl. Fig. 8). First, the specificity of amplicons was verified by cloning and sequencing, including *tbp* (223bp), *psmd14* (150bp) and *app* (247bp). mRNA levels were quantified in cDNA by qPCR with GoTaq qPCR Master Mix (Promega) according to supplier’s instructions in a Mx3000 Real-Time Thermocycler (Stratagene, San Diego, CA, USA). In a 40 cycles PCR reaction, each cycle consisted in 20 s at 94°C, 15 s at 55°C and 15 s at 72°C, followed by a final heating at 95°C, revealing melting curves that confirmed single amplification products. All analyses were performed in triplicate. The expression level of each gene was normalized to *tbp* expression as reference gene using exon-spanning primers to control for genomic DNA contamination since no DNAse treatment of total RNA was included. RT-qPCR assays were analyzed with 2(-ΔΔCt) method ^85^ via MxPro software (Stratagene) and expressed as relative quantity to normalizer ^86^.

### Preparation of protein extracts, Electrophoresis, SDS-PAGE and Western Blot analysis

Cells were washed in ice-cold Phosphate Buffered Saline (PBS) and lysed at 4°C in lysis buffer (50 mM Tris-HCl pH 7.4, 150 mM NaCl, 1 mM EDTA, 1% (vol/vol) Triton X-100) supplemented with a cocktail of protease inhibitors (Sigma-Aldrich). All lysates were cleared by centrifugation at 16,000 x *g* for 20 min at 4°C, and protein concentration was determined with a protein assay dye reagent (Bio-Rad Laboratories, Hercules, CA, USA). Samples with an equivalent amount of protein were boiled for 5 min with Laemmli SDS-PAGE sample buffer, and then analyzed by SDS-PAGE. Proteins were electroblotted onto nitrocellulose membranes, blocked by incubation for 30 min in PBS containing 5% (wt/vol) free-fat dry milk, and incubated sequentially with primary and secondary antibodies, both diluted in blocking solution, for 1 h at room temperature, or overnight at 4°C. Chemiluminescence protein detection was performed using SuperSignal West Pico (Thermo Fisher Scientific). β-Actin was used as an internal loading control.

### In vitro Proteasomal Activity Assay

Proteasome activity was quantitatively assessed in H4 cell extracts using the β5-selective fluorogenic substrate succinyl-leucine-leucine-valine-tyrosine-4-methyl-7-courmarylamide (Suc-LLVY-AMC, Calbiochem, Burlington, MA, USA) using an adapted protocol ^87^. Briefly, cells were left untreated or treated for 4 h with different amounts of CZM (between 2 and 10 μM) or with 10 μM MG132. Cells were lysed in lysis buffer (20 mM Tris-HCl, pH 7.2, 1 mM EDTA, 1 mM dithiothreitol (DTT) and 0.1% (vol/vol) Nonidet P-40) supplemented with a cocktail of protease inhibitors (Sigma-Aldrich). Lysates were cleared by centrifugation at 16,000 x *g* for 20 min at 4°C, and protein concentration was determined with a protein assay dye reagent (Bio-Rad Laboratories, Hercules, CA, USA). Soluble extracts (20 μg) were incubated for 60 min at 37°C in proteasome assay buffer (50 mM Tris-HCl, pH 7.2, 1 mM DTT, 0,5 mM EDTA and 100 μM Suc-LLVY-AMC) all dispensed in triplicate into a 96-well black opaque plate (Nunc, Thermo Fisher Scientific). Proteasome activity was measured by monitoring the production of free AMC using a Synergy HT Multi-detection Microplate Reader (BioTek Instruments, VT, USA) with excitation and emission wavelengths of 360 nm and 460 nm, respectively, and the data were obtained by Gen5 Version 2.09.1 data analysis software. Statistical significance was determined by One-Way ANOVA, followed by Dunnett’s test. Value of P <0,01(**) and P <0,001(***) were regarded as statistically significant and is indicated in the figure.

### Immunofluorescence

Cells grown on glass coverslips were washed with PBS and fixed in 4% (vol/vol) paraformaldehyde for 30 min at room temperature. After fixation, cells were washed in PBS and permeabilized with 0.2% (vol/vol) Triton X-100 in PBS for 10 min at room temperature. Cells were incubated with the indicated primary antibodies diluted in immunofluorescence buffer (PBS containing 10% (vol/vol) FBS and 0.1% (wt/vol) Saponin) for 30 min at 37°C. Coverslips were washed in PBS and incubated with the corresponding Alexa-conjugated secondary antibody diluted in immunofluorescence buffer for 30 min at 37°C. For nuclei staining, cells were washed with PBS and incubated for 10 min at room temperature with 0.1 mg/ml DAPI. After the final wash, coverslips were mounted onto glass slides with Fluoromount-G (SouthernBiotech, Birmingham, AL, USA).

### Fluorescence Microscopy

Images of fixed cells were acquired by using a TCS SP8 laser-scanning confocal microscope (Leica Microsystems, Wetzlar, Germany) equipped with a 63X oil immersion objective (1.4 NA), 405 nm, 488 nm and 561 nm laser lines, with Photomultiplier (PMT), a hybrid detector system (Leica HyD) and the Leica Application Suite LAS X software. For quantification of fluorescent signals, 8-bit images were acquired under identical settings avoiding signal saturation and corrected for background signal on each image. The corrected fluorescent signal in each cell of each image was used in Image J (version 1.44o; Wayne Rasband, NIH, http://imagej.nih.gov) to determine the total integrated pixel intensity per cell area. Colocalization analyses were performed with sets of immunofluorescence images (Z-stack, with 0.2 µm intervals) of the same cells for each marker. Quantification of the acquired images was performed with the ICY software (Quantitative Image Analysis Unit, Institut Pasteur, http://icy.bioimageanalysis.org/) using protocols plugin to create a pipeline to analyze the images in batch, active contours plugin was used to perform the cell segmentation, hk-means plugin was used for threshold detection, wavelet spot detector plugin was used for spot detection and colocalization studio plugin for colocalization analysis.

For live cell imaging assays, H4 cells were grown in glass bottom culture dishes (MatTek Corporation, Ashland, MA, USA) and transiently transfected for 48-h with KDELR-VSVG-YFP, using TransIT-LT1 Transfection Reagent (Mirus Bio LLC, Madison, Wi, USA) according to the manufacturer’s protocol. Before the live cell imaging assay, the culture medium was replaced with phenol red-free DMEM supplemented with HEPES (10mM, pH 7.4), and the cells were treated with CZM (10μM) at 32°C in a controlled temperature chamber in the TCS SP8 laser-scanning confocal microscope. Cells were kept at 32°C to allow KDELR-VSVG-YFP localization on the Golgi, followed by a shift in temperature to 40°C. Imaging was done with a 63X oil immersion objective (1.4 NA), running the Leica Application Suite LAS X software, acquiring 8-bit images at 1-min interval for 15 min at 40°C (488 laser for excitation; HyD: 510-550 nm; 1024×1024 pixels; frame average 1). Quantification of the acquired images was performed with the MetaMorph Software version 7.0.

### 3D Golgi Reconstruction and Golgi Volume and Area Measurements

H4 and HeLa cells stably expressing KDELR1-GFP, under the specified conditions, were fixed and immunostained with anti-GM130 or anti-Giantin, respectively. The immunofluorescence protocol was performed as described above to visualize the Golgi structure. For Golgi volume and area measurements, Z-stack (250 nm) fluorescence images were acquired by using a TCS SP8 laser-scanning confocal microscope (Leica Microsystems, Wetzlar, Germany) equipped with a 63X oil immersion objective (1.4 NA) running the Leica Application Suite LAS X software. Images were then processed with ImageJ software version FIJI to remove the background by using threshold. The thresholded images were then visualized using the “3D Viewer” plugin. The Golgi volume was quantified using ImageJ software version FIJI by setting a thresholded region to select only the Golgi fluorescence. Then, the Golgi structures were separated by ROI (Regions of Interest) and the individual Golgi volume was measured with the plugin “Voxel Counting”. The Golgi volume in μm^3^ was determined by the number of voxels contained in the stack of images (voxel 0.2x0.2x0.25 nm). The Golgi Area was quantified using ICY software and the Golgi marker GM130 was used to determine the ROI. To separate the specific signal from background “k means threshold” plugin was used, using the same threshold level to all images. Data analysis was performed using GraphPad Prism 6 (GraphPad Software, La Jolla, CA, USA) and the results represented in graphs depicting the mean ± SEM of at least 20 cells. Statistical significance of the data was determined with Student’s T-test. The value of P <0,001(***) was regarded as statistically significant and is indicated in the respective figures.

### Densitometric Quantification and Statistical Analysis

The amount of immunoblot signal was estimated using Image J software version 1.48v (Wayne Rasband, NIH, http://imagej.nih.gov). For each condition, protein bands were quantified from at least three independent experiments in order to ensure adequate statistical power. Data analysis was performed using Microsoft Excel 2013 for Windows (Redmond, WA, USA) or GraphPad Prism 6. Results are represented in graphs depicting the mean ± standard deviation. Statistical significance of data comparisons from two groups comparisons was determined with Student’s T-test for parametric data. Values of P <0.05 (*), P <0.01 (**), P <0.001 (***) were regarded as statistically significant and are indicated in the figures. Statistical significance of data from many groups was analyzed using One-Way ANOVA, followed by Tukey’s test in order to evaluate pair-wise comparisons. The value of P <0.05 was regarded as statistically significant and indicated in the figure in different letters above bars mean.

## Author Contributions

Conceptualization, B.H.A., R.F.A. and B.P.V.; Experimental design, B.H.A., R.F.A., M.G.A., C.J., and B.P.V.; Execution of experiments, B.H.A., R.F.A., B.S., C.K., V.G.E., A.M.E., C.T.C., Data analysis, B.H.A., C.K., G.A.E., V.G.E., C.Y., H.S., A.M.E., C.T.C. Reagents, Materials, and Analysis tools, V.G.E., H.S., B.S., A.S., K.G., M.G.A., C.J., H.R.T., R.F.A. and B.P.V. writing—original draft preparation, B.H.A., R.F.A. and B.P.V.; writing—review and editing, B.H.A., B.S., A.S., K.B., M.G.A., C.J., R.F.A., H.R.T and B.P.V.; funding acquisition, B.P.V., R.F.A and H.R.T.

## Funding Statement

This work was funded by Fondo Nacional de Desarrollo Científico y Tecnológico of Chile (FONDECYT; http://www.conicyt.cl/fondecyt) No. 1171649 to B.P.V. & No. 11150532 to R.F.A.; Associative Investigation Program (PIA; https://www.conicyt.cl/pia) including No. ACT-172066 to B.P.V. &No. AFB-170005 to B.P.V.; Academy Insertion Program (PAI; https://www.conicyt.cl/pai) No. 79150075 to R.F.A.; Fondo de Equipamiento Científico y Tecnológico of Chile (FONDEQUIP; http://www.conicyt.cl/fondequip) No. EQM150118 to B.P.V.; Cooperation International Programme (CONICYT-RCUK; https://www.conicyt.cl/pci) No. DPI20140068 to B.P.V.; B.H.A, G.A.E., and V.G.E. were supported by National Ph.D Fellowships No. 21130315, 201110746 and 21130511 respectively (CONICYT; https://www.conicyt.cl/becasconicyt); Programa de Mejoramiento de la Calidad y la Equidad de la Educación superior, MECESUP AUS1203 and Vicerrectoría de Investigación de la Universidad Austral de Chile No. D #2015-02 to B.H.A., D #2013-07 to A.E.G. and D #2015-05 to V.G.E. The funders had no role in study design, data collection and analysis, decision to publish, or preparation of the manuscript.

## Acknowledgments

We thank Gonzalo Astroza (Universidad Austral de Chile) and Ellis Jaffray (GRE, University of Dundee) for technical assistance. Dr. Raymond Deshaies (Division of Biology & Biological Engineering, California Institute of Technology) & Dr. Yuyong Ma (Department of Chemistry and Biochemistry, University of California) for proving Capzimin for this study.

## Conflicts of Interest

The authors declare no conflict of interest.

## Abbreviations

APP: Amyloid Precursor Protein
CZM: Capzimin
DUB: Deubiquitinating enzyme
EBSS: Earle’s balanced salt solution
EGFP: Enhanced Green Fluorescent Protein
ER: Endoplasmic Reticulum
FBS: Fetal bovine serum
GFP: Green Fluorescent Protein
GM130: Golgi matrix protein GM130
HCS: High-Content siRNA Screening
ILV: intraluminal vesicle
K: Lysine residue
KDELR1: KDEL (Lys-Asp-Glu-Leu) Receptor 1
KD: Knock-down
LC3B: Microtubule-associated protein 1 light chain 3B
mTORC: mammalian target of rapamycin complex
MVB: multivesicular body
NT: Non-Target
PBS: Phosphate Buffered Saline
PCR: Polymerase Chain Reaction
PTM: Post-translational Modification
RP: 19S regulatory particle
RT-qPCR: Quantitative reverse transcription PCR
SENP: SUMO-specific protease
siRNA: Small interfering RNA
Ub: Ubiquitin
UBD: Ub-binding domain
ULK1: unc-51-like kinase 1
VSVG: Vesicular Stomatitis Virus G
YFP: Yellow Fluorescent Protein

## Supplementary Figures Legends

**Supplementary Figure 1.**
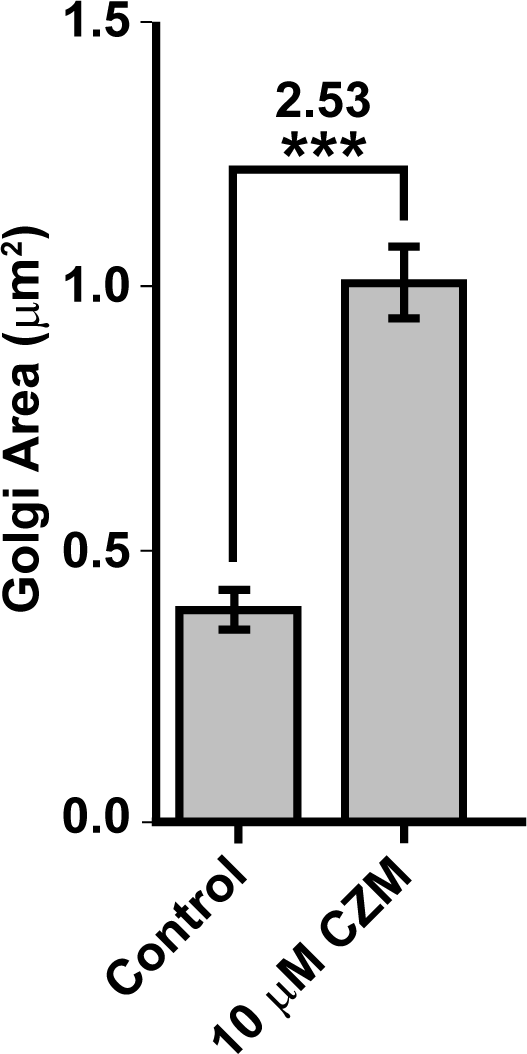
The PSMD14 DUB inhibitor CZM increases the Golgi apparatus area. Immunofluorescence microscopy analysis of the Golgi area in parental H4 cells treated for 4 h either with the vehicle (DMSO; Control) or CZM. The Golgi marker GM130 was used to determine the region of interest in each condition. Statistical significance was determined by Student’s t-test. Bars represent the mean ± SEM (n =43 cells). ***P <0.001.

**Supplementary Figure 2.**
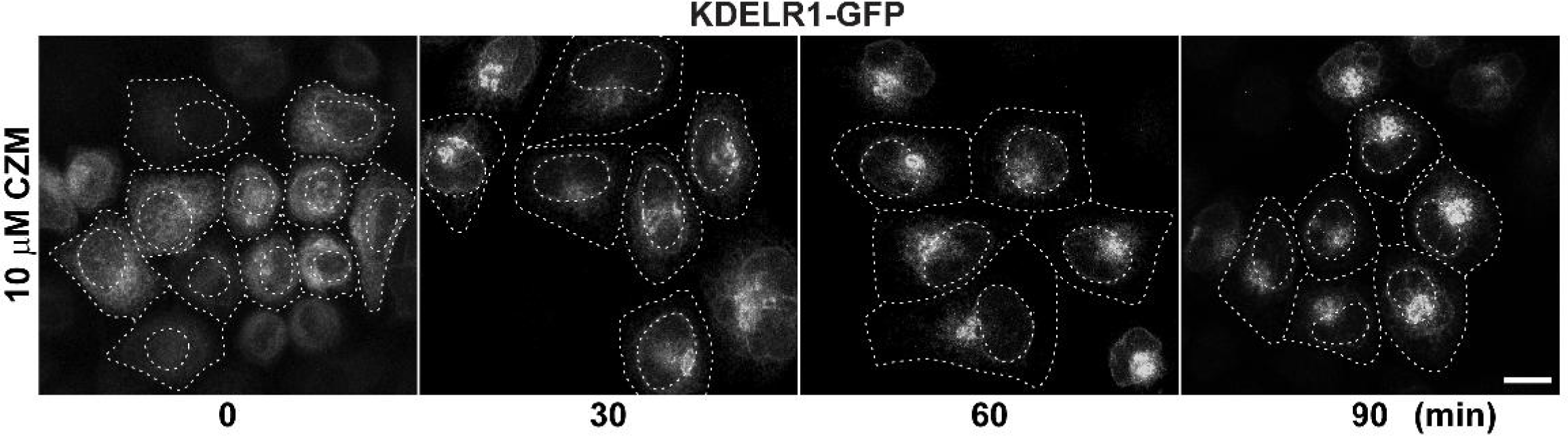
CZM causes the accumulation of KDELR1-GFP at the Golgi apparatus. HeLa cells expressing KDELR1-GFP were either left untreated or treated with CZM for 30, 60 or 90 min. Cells were fixed and representative confocal images were acquired.

**Supplementary Figure 3.**
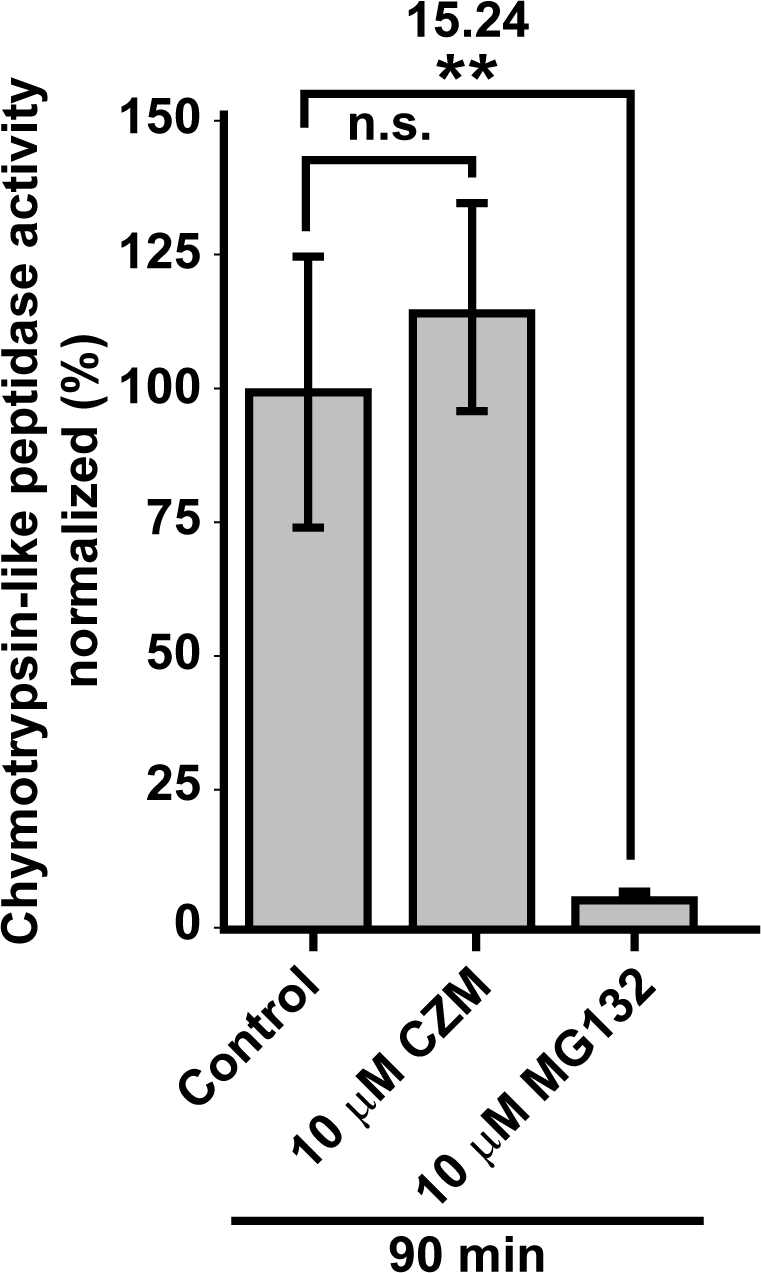
Effect of CZM on proteasome activity. Parental H4 cells were treated either with the vehicle (DMSO; Control), CZM or MG132, for 90 min. Protein extracts were used to measure *in vitro* the Chymotrypsin-like peptidase activity of the proteasome. The enzymatic activity was quantified according to the cleavage of the fluorogenic substrate Suc-LLVY-AMC to AMC, and normalized to that of control cells. The statistical significance was determined by One-Way ANOVA, followed by Tukey’s test. Bars represent the mean ± SD of biological replicates (n=3). **P <0.01; n.s., not significant.

**Supplementary Figure 4.**
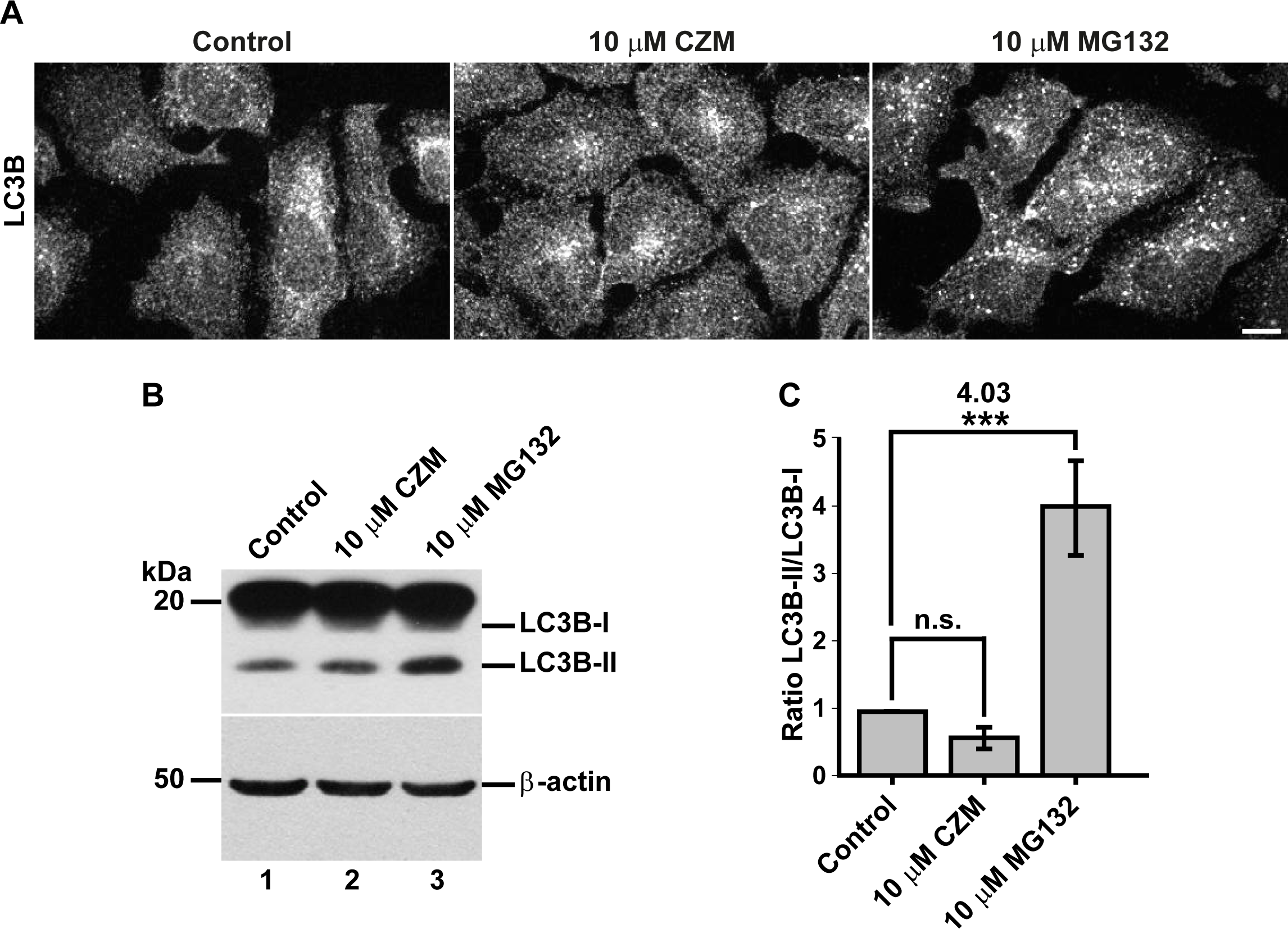
Effect of CZM and MG132 on basal macroautophagy. **(A)** Immunofluorescence microscopy analysis of the subcellular localization of LC3 in parental H4 cells treated with either with the vehicle (DMSO; Control), CZM for 4 h or MG132 for 6 h. Cells were fixed, permeabilized and stained with a rabbit polyclonal antibody to LC3B followed by Alexa-594-conjugated donkey anti-Rabbit IgG. Scale bar 10 µm. **(B)** Parental H4 cells were treated as in (A) and protein extracts were analyzed by western blot with a polyclonal antibody to LC3B. Monoclonal antibody to β-actin (clone BA3R) was used as a loading control. The position of molecular mass markers is indicated on the left. **(C)** Ratio LC3B-II/LC3B-I ratio as shown in (B). The statistical significance was determined by One-Way ANOVA, followed by Tukey’s test. Bars represent the mean ± SD of biological replicates (n=3). ***P <0.001; n.s., not significant.

**Supplementary Figure 5.**
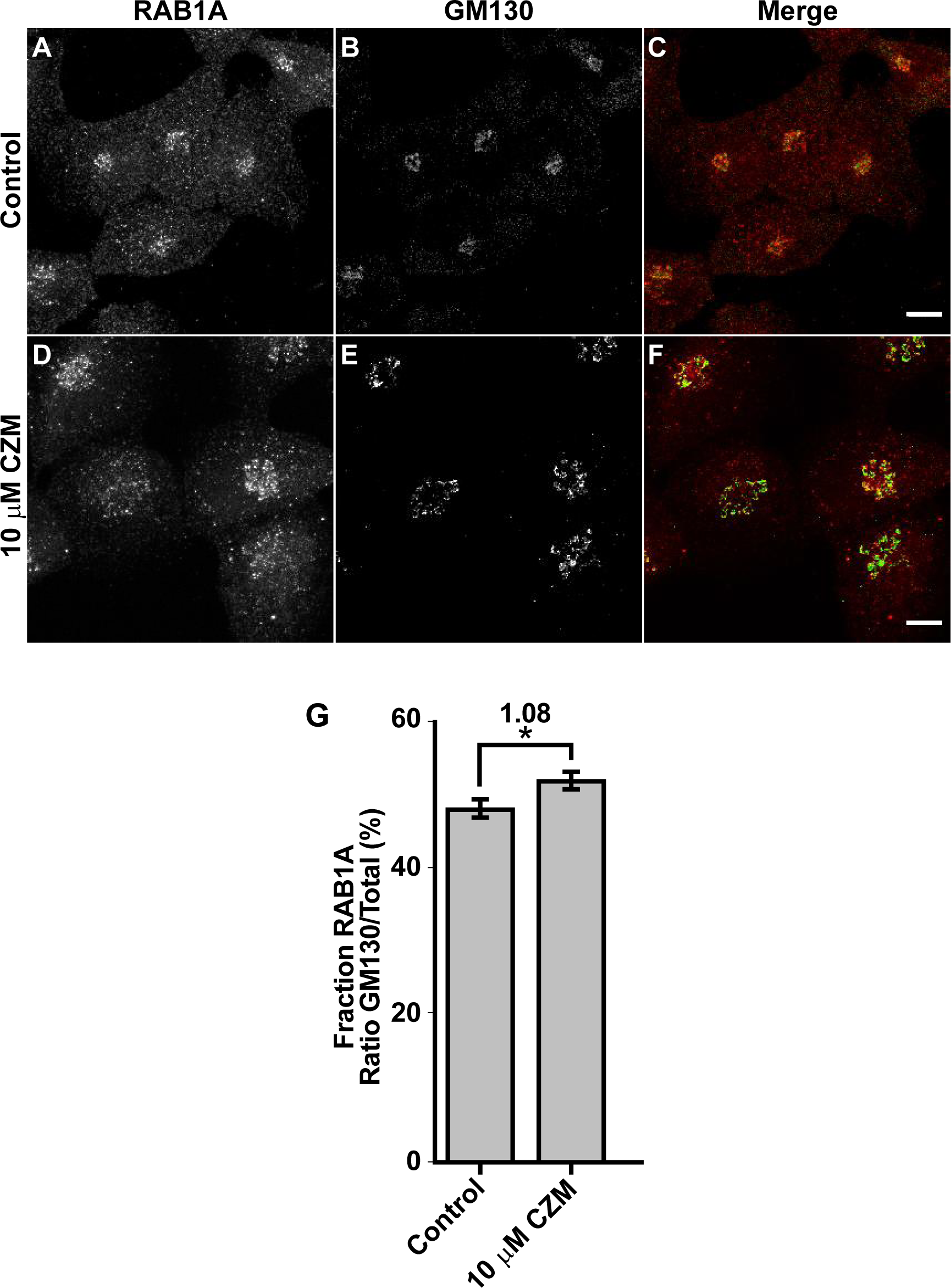
Distribution of RAB1A upon CZM treatment. Immunofluorescence analysis of endogenous RAB1A in H4 parental cells treated either with vehicle (DMSO; Control) **(A-C)** or CZM for 4 h **(D-F)**. Cells were fixed, permeabilized, and double stained with a rabbit monoclonal antibody to RAB1A (clone D3X9S) (A and D) and a mouse monoclonal antibody to GM130 (clone35/GM130) (B and E), followed by Alexa-594-conjugated donkey anti-Rabbit IgG and Alexa-488-conjugated donkey anti-Mouse IgG. Merging of the images generated the third picture (C and F). Scale bar, 10 μm. **(G)** Quantitative analysis of the fraction of RAB1A colocalizing with GM130 under CZM treatment and compared to control cells. The statistical significance was determined by Student’s t-test. Bars represent the mean ± SEM of the fluorescent signal per cell area (n=173 cells). *P< 0.05.

**Supplementary Figure 6.**
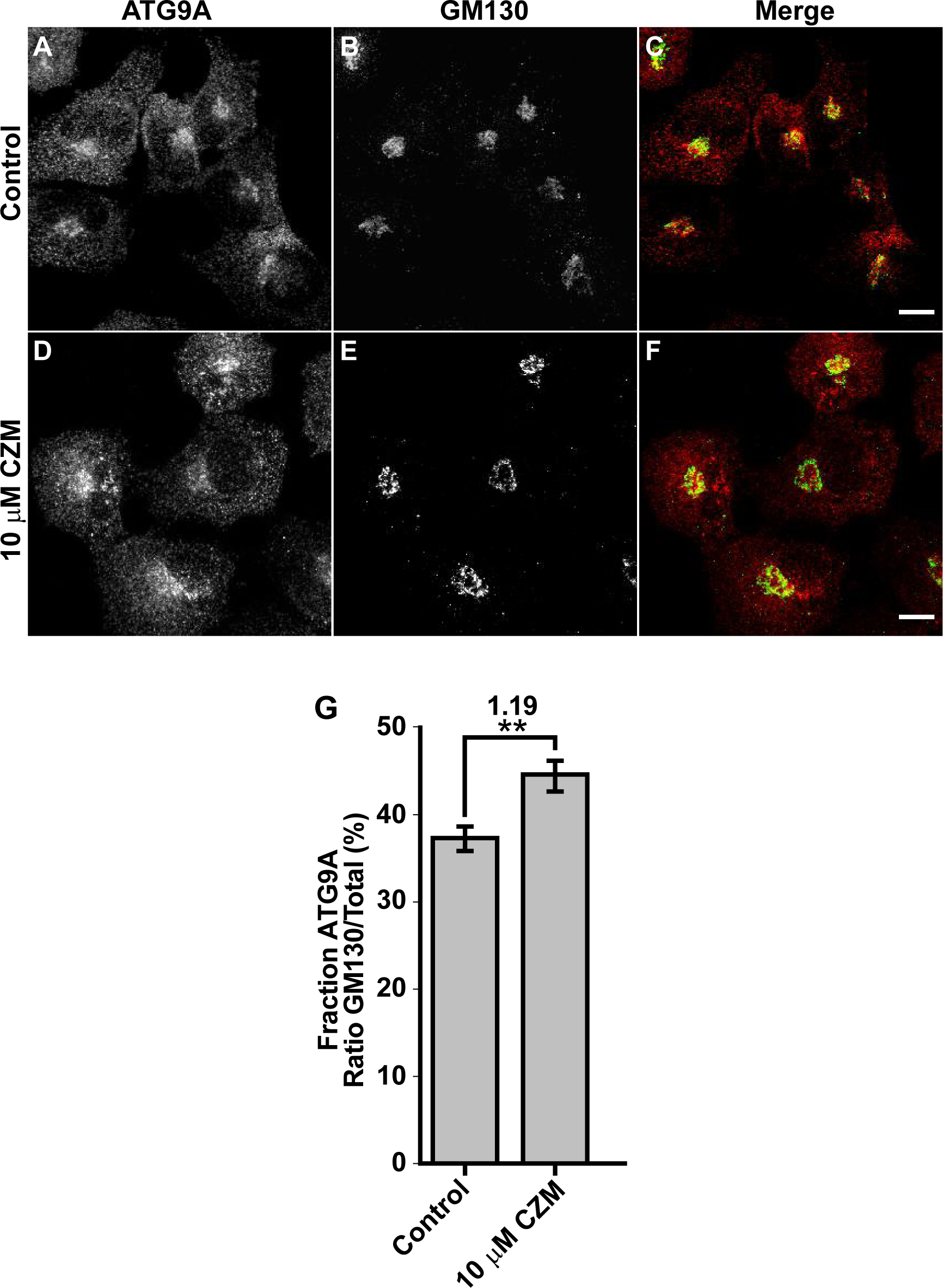
ATG9A is distributed in the swollen Golgi apparatus upon CZM treatment. Immunofluorescence analysis of endogenous ATG9A in H4 parental cells treated either with the vehicle (DMSO; Control) **(A-C)** or CZM for 4 h **(D-F)**. Cells were fixed, permeabilized, and double stained with a rabbit monoclonal antibody to ATG9A (clone EPR2450(2)) (A and D) and a mouse monoclonal antibody to GM130 (clone35/GM130) (B and E), followed by Alexa-594-conjugated donkey anti-Rabbit IgG and Alexa-488-conjugated donkey anti-Mouse IgG. Merging of the images generated the third picture (C and F). Scale bar, 10 μm. **(G)** Quantitative analysis of the fraction of ATG9A colocalizing with GM130 under CZM treatment and compared to control cells. The statistical significance was determined by Student’s t-test. Bars represent the mean ± SEM of the fluorescent signal per cell area (n=93 cells). **P <0.01.

**Supplementary Figure 7.**
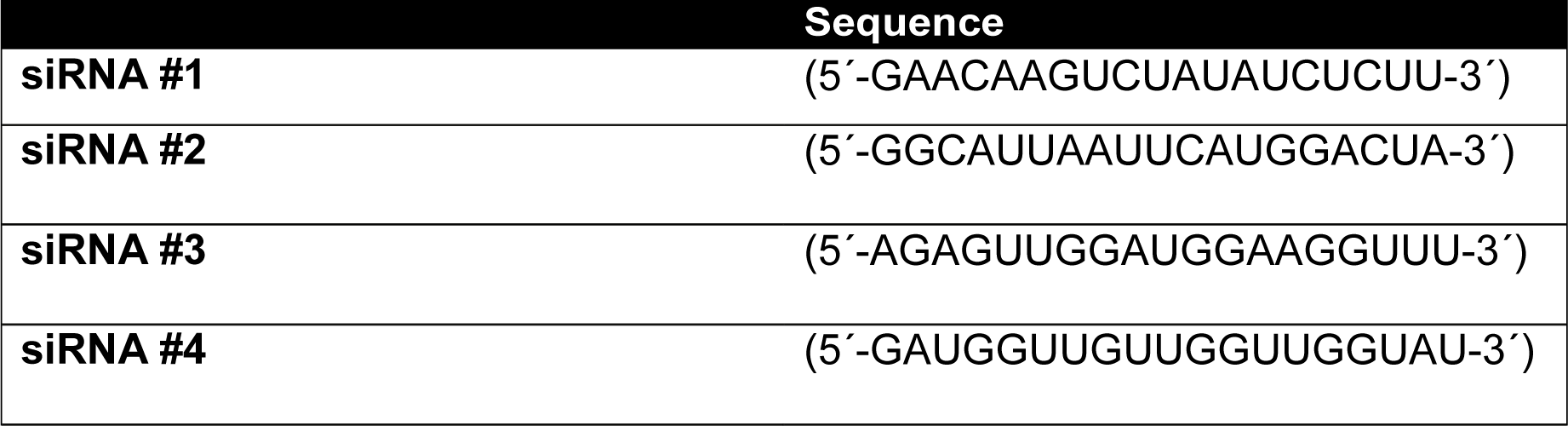
siRNA sequences directed against human PSMD14 used for Validation Stage.

**Supplementary Figure 8.**
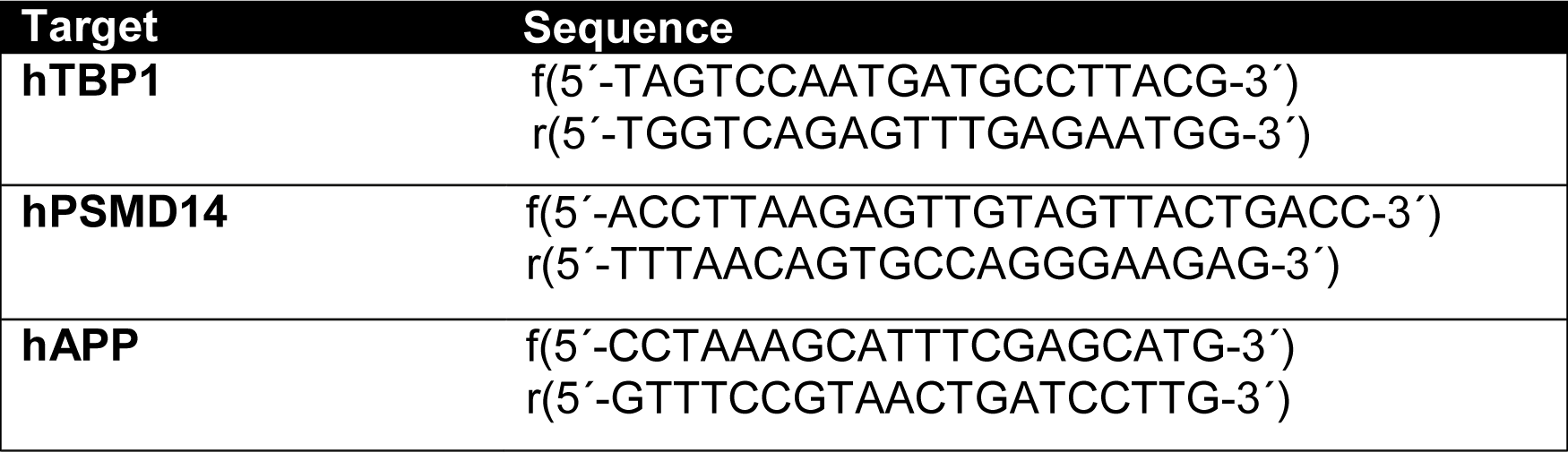
Primer pairs sequences used for RT-qPCR.

